# Pre-mature senescence in the oldest leaves of low nitrate-grown *Atxdh1* mutant uncovers a role for purine catabolism in plant nitrogen metabolism

**DOI:** 10.1101/233569

**Authors:** Aigerim Soltabayeva, Sudhakar Srivastava, Assylay Kurmanbayeva, Aizat Bekturova, Robert Fluhr, Moshe Sagi

## Abstract

The nitrogen rich ureides allantoin and allantoate, are known to play a role in nitrogen delivery in *Leguminosae*, in addition to their role as reactive oxygen species scavengers. However, their role as a nitrogen source in non-legume plants has not been shown. *Xanthine dehydrogenase1* (AtXDH1) activity is a catalytic bottleneck step in purine catabolism. *Atxdh1* mutant exhibited early leaf senescence, lower soluble protein and organic-N levels as compared to wild-type (WT) older leaves when grown with 1 mM nitrate, whereas under 5mM, mutant plants were comparable to WT. Similar nitrate-dependent senescence phenotypes were evident in the older leaves of allantoinase (*Ataln*) and allantoate amidohydrolase (*Ataah*) mutants, impaired in further downstream steps of purine catabolism. Importantly, under low nitrate conditions, xanthine was accumulated in older leaves of *Atxdh1*, whereas allantoin in both older and younger leaves of *Ataln* but not in WT leaves, indicating remobilization of xanthine degraded products from older to younger leaves. Supporting this notion, ureide transporters *UPS1, UPS2* and *UPS5* were enhanced in older leaves of 1 mM nitrate-fed WT as compared to 5 mM. Enhanced AtXDH, AtAAH and purine catabolic transcripts, were detected in WT grown in low nitrate, indicating regulation at protein and transcript levels. Higher nitrate reductase activity in *Atxdh1* than WT leaves, indicates their need for nitrate assimilation products. It is further demonstrated that the absence of remobilized purine-degraded N from older leaves is the cause for senescence symptoms, a result of higher chloroplastic protein degradation in older leaves of nitrate starved *Atxdh1* plants.

**Summary:** The absence of remobilized purine-degraded N from older to the young growing leaves is the cause for senescence symptoms, a result of higher chloroplastic protein degradation in older leaves of nitrate starved *Atxdh1* plants.

## INTRODUCTION

In plants, the degradation of purine compounds starts with the conversion of adenosine monophosphate (AMP) to inosine monophosphate (IMP) by AMP deaminase (AMPD, EC 3.5.4.6), which leads, by multiple pathways, to the production of oxypurines such as xanthine and hypoxanthine (Yoshino *et al.*, 1979; Xu *et al.*, 2005; Zrenner *et al.*, 2006; Sabina *et al.*, 2007). In the degradation pathway, xanthine is initially oxidized to urate, that is further converted by urate oxidase (UO, EC 1.7.3.3) and a transthyretin-like protein to allantoin, the main end product in most mammals (Zrenner *et al.*, 2006; Reumann *et al.*, 2007; Werner and Witte, 2011; Hauck *et al.*, 2014). Conversely, plants possess a set of enzymes which further break down allantoin to ureidoglycolate, catalyzed by allantoinase (ALN, EC 3.5.2.5), allantoate amidinohydrolase (AAH, EC 3.5.3.9.) and ureidoglycine aminohyrolase (UGlyAH, EC 3.5.3-.) (Werner *et al.*, 2010, 2013). The ureidoglycolate amidohydrolase (UAH, EC 3.5.1.116.) converts ureidoglycolate to the basic metabolic building blocks, glyoxylate and ammoinium (Werner *et al.*, 2013). The release of four ammonium molecules, that essentially should be reassimilated, parallel the sequential hydrolysis of purines (Werner *et al.*, 2010).

Environmental stimuli can induce premature leaf senescence (Miller et al., 1999; Munne-Bosch and Alegre, 2004; Pageau et al., 2006; Lim et al., 2007). Natural leaf senescence occurs in a coordinated manner (Lim *et al.*, 2003); it starts from inhibition of leaf expansion (Diaz *et al.*, 2005), followed by induction of metabolic changes that result in nutrient degradation and remobilization (Pate, 1980; Simpson *et al.*, 1983). This strategy of recycling endogenous nutrients from the senescing leaves can be used by plants to maintain growth of younger leaves and reproductive organs under nutrient-limiting stress (Aerts, 1990; Buchanan-Wollaston and Ainsworth, 1997; Hortensteiner and Feller, 2002; Eckhardt et al., 2004). In the senescing leaves most nitrogen is contained in proteins (Masclaux-Daubresse *et al.*, 2010), but the nucleobases such as purines are also rich in nitrogen (Thomas *et al.*, 1980; Atkins *et al.*, 1982; Schubert, 1986; Brychkova *et al.*, 2015) and thus may be a source for nitrogen (N) recycling. In legumes the N fixed in the form of ureides was indeed shown to be translocated from the nodules, where the ureides are synthesized *de novo*, to the aerial plant tissues where they are degraded and used as N source (Stebbins and Polacco, 1995; Smith and Atkins, 2002b; Todd et al., 2006). A role for nucleic acid/purine degradation products in plant nitrogen metabolism was presented also in non N-fixing legumes. The recycling of nucleic acids in tissues undergoing stress-induced pre-mature senescence was considered as a possible source for ureides increase in non-nodulated common bean plants (Alamillo *et al.*, 2010). Impressively, the high levels of ureides evident in shoot and leaves of non-nodulated *Phaseolus vulgaris* plants fertilized with nitrate was suggested to be the result of remobilized N from senescent leaves to be employed for new growing tissue (Díaz-Leal *et al.*, 2012).

Interestingly, uric acid, allantoin, and allantoate, the products of purine degradation, were shown to be able to serve as sole nitrogen sources during the growth of Arabidopsis plants when supplemented externally to the growth medium (Desimone *et al.*, 2002; Todd and Polacco, 2004; Nakagawa *et al.*, 2007). However, it was claimed that Arabidopsis plants grown on allantoin did not seem to perceive it as a good sole N source (Werner et al., 2008) exhibiting a reduced growth (Desimone *et al.*, 2002).

We have previously demonstrated that aging and extended darkness activates the purine catabolism pathway to accumulate ureides, which act as antioxidants against oxidative stress, and delay pre-mature and natural senescence in Arabidopsis leaves (Brychkova *et al.*, 2008). Yet, the role of the senescence-induced purine degradation products such as ureides (Brychkova *et al.*, 2008) has not been fully examined in plants and no functional analysis employing relevant mutants has yet proved that purine degradation products has a role in nitrogen metabolism. The fact that leaf senescence is paralleled by a decrease in RNA (Crafts-Brandner *et al.*, 1996; Crafts Brandner *et al.*, 1998) and an increase in the level of uriedes, whereas enzymes/transcripts of genes involved in the purine degradation pathway are up-regulated during leaf senescence (Brychkova *et al.*, 2008), gives more than a clue as to the involvement of RNA degraded products in N metabolism (Werner and Witte, 2011, Have et al., 2016).

To examine the role of purine degraded metabolites as an important nitrogen source in Arabidopsis plant development, the knockout mutants *Atxdh1*, *Ataln* and *Ataah* defective in Xanthine dehydrogenase (*XDH1*), Allantoinase (*ALN*) and Allantoate amidohydrolase (*AAH*) expression respectively, were studied under sufficient and limited nitrogen conditions. Under nitrogen deficient conditions the purine degradation pathway is activated on transcript and protein levels to provide an additional source of nitrogen from older senescent leaves to the young leaves. Growth of the purine catabolism mutants under nitrate limitation resulted in premature senescence symptoms in old leaves, but not in those of WT plants. In contrast, sufficient supply of nitrate resulted in almost full disappearance of the premature senescence symptoms, with parallel enhancement of the organic nitrogen and soluble protein content in the mutant older leaves. This was achieved by higher nitrate reductase activity in *Atxdh1* than WT leaves. Further, it was demonstrated that the absence of remobilized purine-degraded N from older leaves is the cause for senescence symptoms, a result of higher degradation of chloroplastic proteins, such as Rubisco large subunit and D1, the component of the reaction center of PSII, in older leaves of nitrate starved *Atxdh1* plants.

## RESULTS

### High nitrate supplementation prevents senescence symptoms in *Atxdh1* older leaves

Previously it was shown that a mutation in *Atxdh1*, a key enzyme in the purine catabolism procees, confers early leaf senescence (Brychkova et al., 2008). To examine the role of purine degraded metabolites as an important endogenous nitrogen source, the effect of sufficient and limited nitrogen supplementation to the knockout mutants *Atxdh1*, *Ataln* and *Ataah* defective in *XDH1*, *ALN* and AAH, respectively was studied with 25 days old plants. Total chlorophyll levels in *Atxdh1*, *Ataln* and *Ataah* mutants old leaves supplemented with low nitrate (1 mM) were lower than in wild type (WT) old leaves, whereas no difference was noticed in the young leaves. Importantly, increasing nitrate levels in the growth medium (5 mM) enhanced the total chlorophyll level in the mutants old leaves (Fig. 1AB, Supplementary Fig. S1). Additionally, the senescence marker Cys protease senescence-associated gene 12 [*SAG12* (Gepstein *et al.*, 2003)], the chlorophyll-degradation gene *ACD2* [accelerated cell death2 (Tanaka *et al.*, 2003)] and stay-green protein1 [*SNG1* (Park et al., 2007)] were significantly upregulated in the old leaves of the mutants supplemented with the lowest nitrate level as compared to WT, but not in mutants old leaves in plants grown with high nitrate (Fig. 1C, Supplementary Fig. S1 C, D). These results indicate that limited nitrate supply resulted in enhanced chlorophyll degradation in mutants impaired in purine catabolism genes. This is likely due to a shortage of endogenous nitrogen sources in the mutant lines that could be repaired by higher nitrate supplementation.

**Figure 1.**
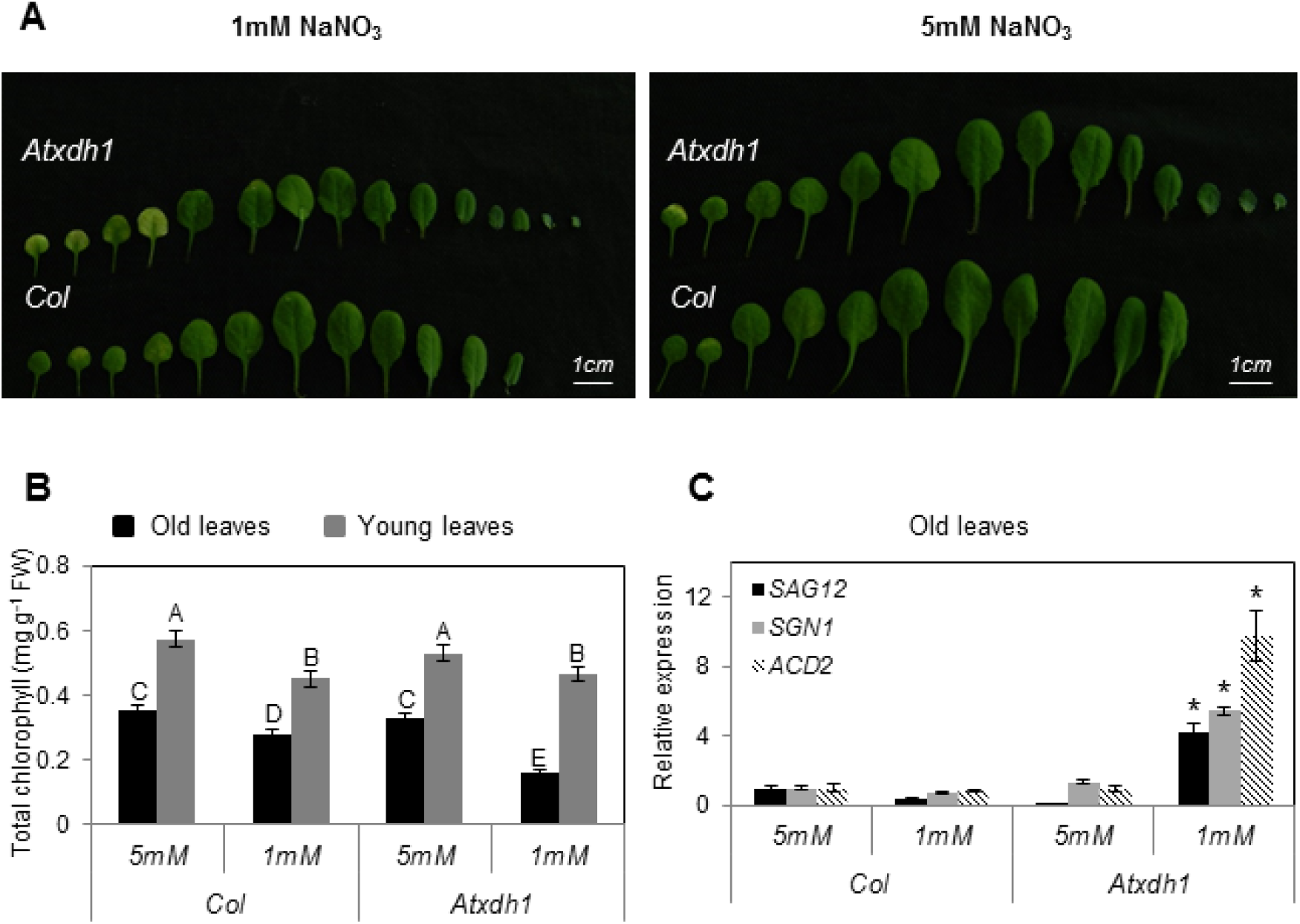
Effects of different nitrate levels, supplemented to the growth medium of WT (Col) and *Atxdh1* mutant, on senescence symptoms in leaves. Leaf appearance (A) from left to right is young to old where the first and last 4 leaves are designated ‘young’ and ‘old’, respectively. Total chlorophyll (B) content. Relative expression (C) of senescence marker transcripts in old leaves of *SAG12* (Suppressor of overexpression of Cys protease senescence-associated gene 12), *ACD2* (accelerated cell death2), *SGN1* (stay-green protein 1) (At5G45890, At4G37000, At4G22920, respectively). The data of chlorophyll content represent the mean obtained from a representative experiment from six independent biological replications. Values denoted by different letters are significantly different (Tukey-Kramer HSD test, P < 0.05). The expression of each transcript was compared with the young leaves of WT in 5 mM nitrate treatment after normalization to *EF-1α* transcript (At5g60390). Values marked with asterisk denotes significant difference (T-test, n=3, P < 0.05) between treatment and genotypes for each transcript and the data represent the mean obtained from 3 independent experiments. *Atxdh1* are SALK_148366 and GABI 049004 T-DNA mutants

### *Atxdh1* mutation confers lower organic nitrogen and soluble protein levels, but higher RNA than in WT old leaves grown under N deficiency

Levels of total nitrogen in its various forms, are key physiological indicator of plant health. Examination of nitrogen (N) levels in the young and old leaves grown under low (1mM) and high (5mM) nitrate, revealed a total N decrease in WT and *Atxdh1* leaves with decreasing nitrate application. However, total nitrogen (N) level was significantly lower in old leaves of *Atxdh1* mutant as compared to WT (Fig. 2A). Importantly, under the low nitrogen supplementation the organic N level was considerably lower in the old and young leaves of the *Atxdh1* mutant as compared to WT (decrease of 1.03 and 0.83 mmol N per g DW, respectively), whereas, no significant difference was noticed between the leaves of these two genotypes when fed with high nitrate (Fig. 2B). This indicates that under low nitrate conditions the mutation in *XDH1* stimulates the degradation of organic nitrogen and protein in the older leaves (Fig. 2), to supply nitrogen essential for the growth of the younger leaves. This is consistent with the early senescence phenotype in mutant older leaves and its absence in the younger leaves, albeit the organic N in the latter was lower than in WT, but still significantly higher as compared to the old leaves in *Atxdh1* plant grown under low nitrate supply (Fig. 1, 2).

**Figure 2.**
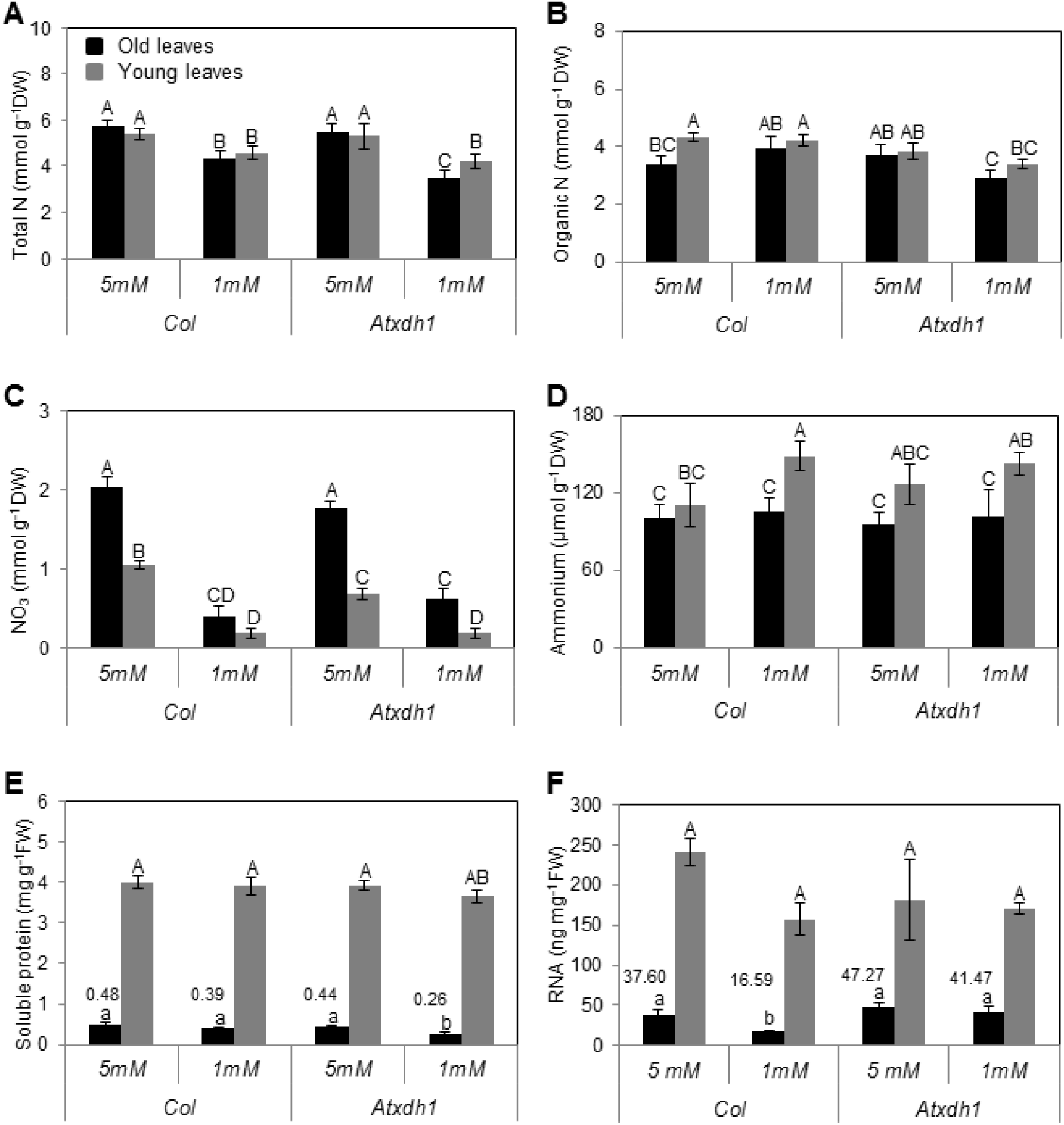
Effects of different nitrate levels on total N (A), total organic N (B), nitrate (C), ammonium (D), soluble protein content (E) and totalRNA content (F) in old and young leaves of WT (Col) and *Atxdh1* mutants. Plants were grown until 25 days old in nitrogen deficient soil supplemented with one-half strength Hoagland nutrient solution supplemented with 1 or 5mM NaNO_3_ as the only N source. The data represent the mean obtained from six independent experiments. The values denoted with different letters are significantly different according to the Tukey-Kramer HSD test, (P < 0.05). Different upper case letters in inserts (A) to (D) indicate differences between treatments. Different capital letters in inserts (E) and (F) indicate differences between treated young leaves. Different lower case letters in inserts (E) and (F) indicate differences between treated old leaves. *Atxdh1* are SALK_148366 and GABI_049004 T-DNA mutants

The high nitrate application resulted in increased nitrate content in the young and old WT and *Atxdh1* leaves, being lower in young leaves of *Atxdh1* mutant as compared to WT plants (Fig. 2C). The significantly lower nitrate accumulation in *Atxdh1* younger leaves is indicative for a higher rate of nitrate assimilation in the mutant younger leaves, to overcome the absence of ammonium, originated from unimpaired purine catabolism and employed for organic N biosynthesis in WT. Indeed, the ammonium content in the *Atxdh1* mutant was similar as in the WT grown under high nitrate conditions (Fig. 2D) indicating that nitrate was assimilated to ammonium, to be incorporated into organic molecules (Somerville and Ogren 1980; Joy 1988; Stitt, 1999; Wang et al., 2003, Coruzzi 2003). Interestingly, total soluble protein content in young leaves was similar in WT and mutant plants being significantly higher than the old leaves of both genotypes. Yet, soluble proteins were significantly lower in old leaves of 1mM nitrate fed mutant plants as compared to WT [decreased by 33% (by 0.13 mg soluble protein per g FW)], the latter having similar soluble protein levels as the 5 mM supplied WT and mutant plants (Fig. 2E). The lowest soluble protein and organic N in old leaves of the mutant fed with low nitrate (Fig. 2B, E) is indicative for remobilization of degraded protein products from these leaves.

The estimation of total RNA level does not represent the whole pool of purine degraded compounds, since there are additional purine pools in plants such as nucleosides and bases (eg. AMP, ADP, ATP, GMP, GDP), purine alkaloids (eg. 3-Methylxanthine, 7-methylxanthosine, Theobromine), coenzymes (e.g. NAD, NADP, FAD, coenzyme A) and adenylosuccinate, as well as isopentenyl adenosine monophosphate and more (Meyer and Wagner, 1986; Smith and Atkins, 2002; Koyama *et al.*, 2003; Lange *et al.*, 2007; Sabina *et al.*, 2007; Ashihara *et al.*, 2008; Agrimi *et al.*, 2012) Yet, the RNA estimation is a good indicator for the process, especially when considering the level of the ureides accumulated in *Ataln* mutants (see below). With this caveat in mind, the significant low RNA level in WT old leaves fed with low nitrate as compared to the RNA levels in WT supplemented with high nitrate or mutant leaves fed with low or high nitrate is indicative for the employment of purine degradation product for N remobilization from these leaves to the upper leaves, where no differences in RNA levels were evident within nitrate treatments or between genotypes (Fig. 2F). Overall, the significant rate of decrease in total N, organic N, and soluble proteins and the higher RNA in the old mutant leaves as compared to WT old leaves in low nitrate fed plants, suggests the neglegible N remobilization from purines and the significant N degradation and remobilization from mutant old leaves proteins.

### Low nitrogen supplementation confers enhancement of ureide transporter transcripts in old leaves of WT

Ureides, the purine degradation products, were shown to be transported from the nodules of legume roots via the xylem (Collier and Tegeder, 2012) to the shoot (Schubert, 1981). By employing orthologue *AtUPS* gene expression in yeast and/or xenopus with other than Arabidopsis promoters it was shown that the AtUPS1 acts as xanthine and allantoin permeases whereas AtUPS2 is an uracil, and AtUPS5 is likely more a xanthine and allantoin permease (Desimone *et al.*, 2002; Schmidt *et al.*, 2004, 2006). Recently, *AtUPS5* was suggested to act as a key component in allantoin transport to the shoots (Schmidt et al., 2006, Lescano, 2016), whereas *AtUPS1* was shown to be significantly up-regulated in adult Arabidopsis shoot as a response to a sudden total nitrogen starvation (Krapp *et al.*, 2011). The lower total RNA level in old leaves of nitrogen starved WT as compared to *Atxdh1* (Fig. 2F), led us to explore the transcript expression of these UPS transporters. Overall, older leaves had higher levels of all transporters compared to younger leaves. The expression of *AtUPS1*, *AtUPS2* and *AtUPS5* was especially high in old WT leaves of plants fed with 1 mM nitrate compared to plants supplemented with 5 mM nitrate or in *Atxdh1* mutant plants (Fig. 3). Importantly, the expression of *AtUPS1* and *AtUPS2* in old and young leaves of low nitrate fed *Ataln* plants was folds higher than in the WT plant, whereas *AtUPS5* in old leaves of *Ataln* was similar as shown in WT supplemented with low nitrate (Fig. 3). These results support the notions of i) the role of *AtUPS1*, *AtUPS2* and *AtUPS5* in mediating ureide transport and ii) ureide remobilization from the old leaves to the young growing leaves of nitrate starved plants. It also suggests that induction of the *UPS* transcripts is sensitive to the flux of catabolic products and these are lacking in the *Atxdh1* mutant.

**Figure 3.**
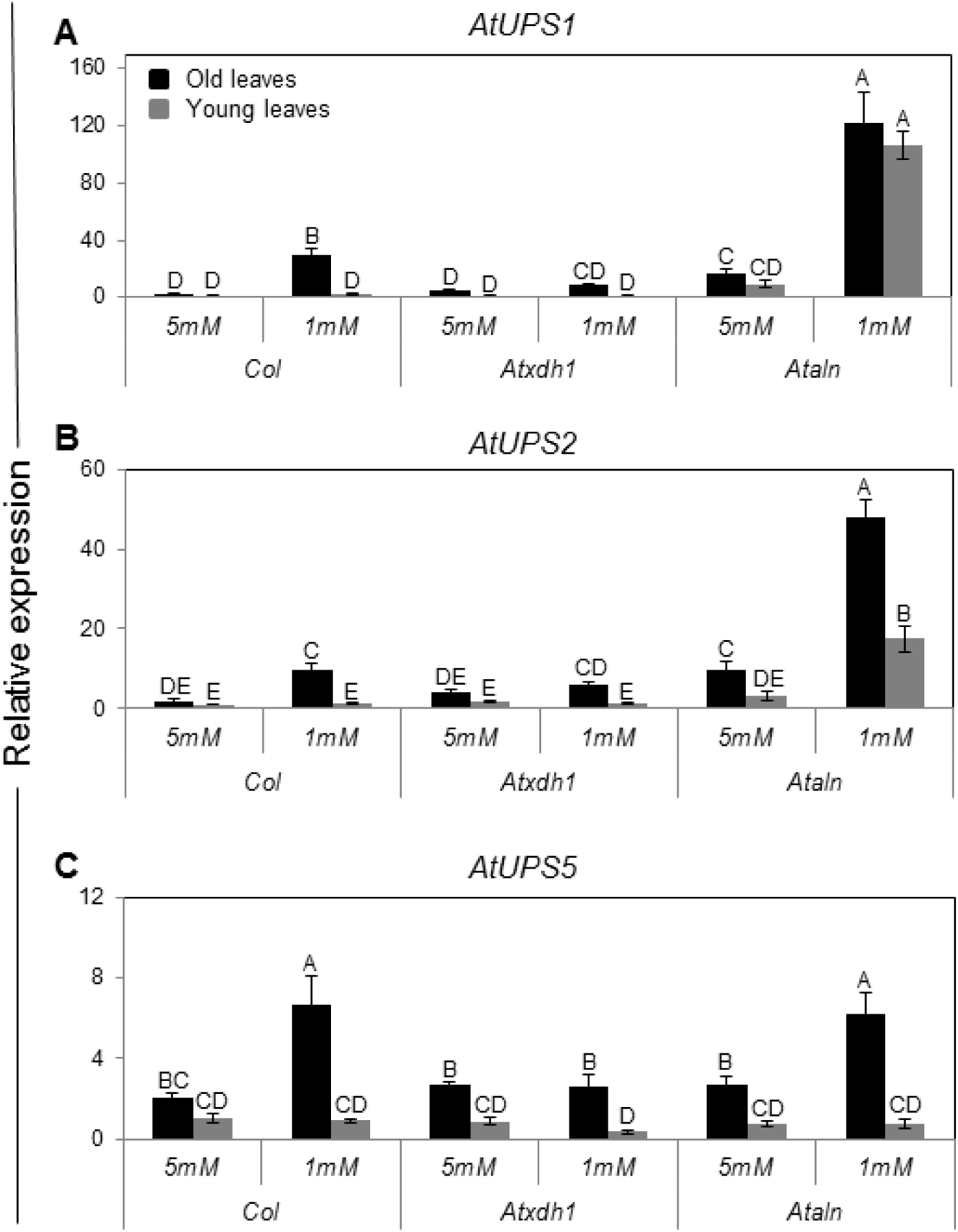
Transcript expression levels of the ureide permeases, *AtUPS1*(A), *AtUPS2* (B) and *AtUPS5* (C), in old and young leaves of WT (Col), *Atxdh1* and *Ataln* mutants in response to nitrate level supplemented to the growth medium. Quantitative analysis of transcripts by real-time RT-PCR was performed using 25 old plants grown on nitrogen deficient soil supplemented with 1 or 5mM NaNO_3_ as the only N source. The expression of each treated line was compared with the young leaves of Col in 5mM nitrate treatment after normalization to *EF-1α* transcript (At5g60390). The data represent the mean obtained from three independent experiments (Tukey-Kramer HSD test, P < 0.05). *Atxdh1* and *Ataln* are SALK_148366 and SALK_013427 T-DNA mutants respectively.

### Xanthine and allantoin catabolism in WT and mutants impaired in purine catabolism

In case of catabolic activity substrates are expected to accumulate within the mutant lines for purine catabolism. It is therefore of interest to examine the possibility of differential accumulation in WT and mutants young and old leaves. Importantly, xanthine accumulated chiefly in the old leaves of *Atxdh1*, was several folds higher than in the young leaves and was significantly higher in old leaves in plants supplemented with low nitrate compared to old leaves of *Atxdh1* fed with high nitrate. The xanthine level in WT old leaves was folds lower than in the mutant (Fig. 4A). The possible stress effect of xanthine toxicity was examined in leaf discs sampled from 6^th^ to 10^th^ rosette leaves (counted from the bottom and being without senescence symptoms) exposed to water (mock) and 1mM xanthine or allantoin for 48 h. Anthocyanin accumulation was used as a sign of stress (Chalker-Scott, 1999; Gould *et al.*, 2002; Schussler *et al.*, 2008). Higher anthocyanin levels were evident in the presence of xanthine as compared to allantoin especially in *Atxdh1* when compared to WT leaf discs (Supplementary Fig. S2). This indicates that xanthine accumulation could contribute to stress and hastening of the senescence phenotype in *Atxdh1* old leaves supplemented with low N (See in Fig. 1) especially in the absence of ureides (Figure 8a in Brychkova et al., 2008). However, this is likely not the case here (Fig. 1), since no senescence symptoms could be noticed in the xanthine treated leaf discs (See in Supplementary Fig. S2). Moreover, the level of xanthine accumulated in the old leaves of *Atxdh1* mutant (Fig. 4A) was far lower than the level shown recently in *Atxdh1* leaves after 5 successive days in dark [calculated as ∼1 μmol g^-1^ FW from Figure 8A in (Schroeder *et al.*, 2017)], where no significant senescence symptoms is claimed to be more in the mutant compared to WT leaves (See Figure 7 in Schroeder et al., 2017). Significantly, only low xanthine levels were evident in the old leaves of *Ataln* and *Ataah* mutants and WT plants (Fig. 4A, Supplementary Fig. S3). Yet, the mutants also displayed enhanced senescence symptoms relative to WT under low nitrate conditions (Fig. 1 and Supplementary Fig. S1). This suggests that the absence of the nitrogen rich allantoin (Fig. 4B) degraded products demanded for the young leaf growth, was substituted by the degraded product of older leaf chloroplastic-proteins as as can be seen by the higher chlorophyll degradation and senescence marker transcripts *SAG12* and *SGN1* in these leaves (Suplementary Fig. S1).

**Figure 4.**
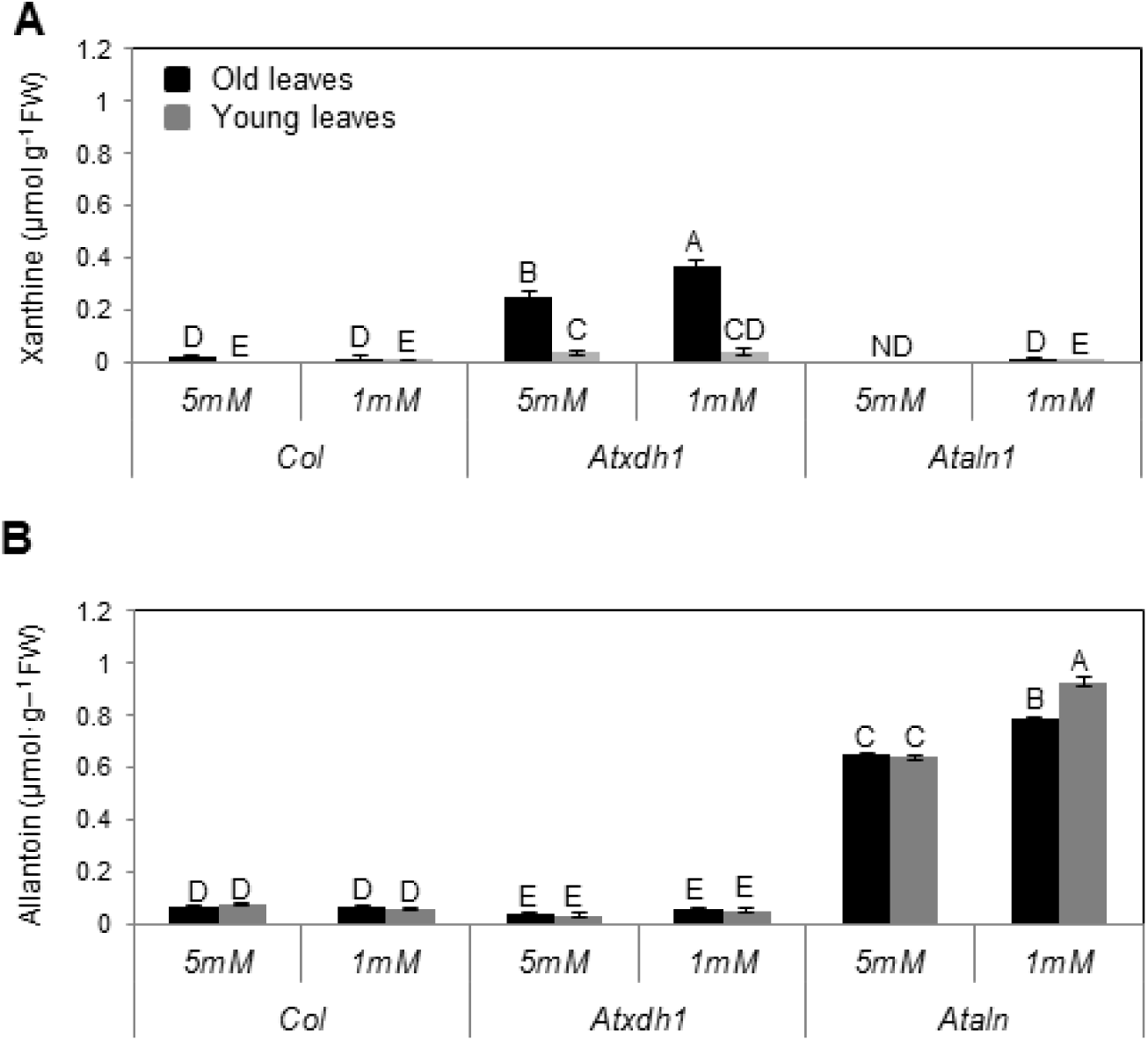
The levels of xanthine (A), allantoin (B) in old and young leaves of WT (Col), *Atxdh1, Ataln* mutants grown in nitrogen deficient soil supplemented with 1 or 5 mM NaNO_3_. ND-not detected. The data represent the mean obtained from one of 5 independent experiments with similar results. The values denoted with different letters are significantly different according to the Tukey-Kramer HSD test, (p< 0.05). Different upper case letters indicate differences between mutants and WT plants. *Atxdh1* are SALK_148366 and GABI_049004, whereas *Ataln* are SALK_13427 and SALK_46783 T-DNA mutants.

Xanthine accumulation in *Atxdh1* mutant reports higher purine catabolic activity (Ma *et al.*, 2016). The several folds higher xanthine levels accumulated in *Atxdh1* old leaves, being the highest in the low nitrate fed plants is thus indicative for the high demand for purine degradation products blocked from being further catabolized and remobilized for young leaf growth (Fig 4A). The low xanthine level in WT old leaves is a result of xanthine degradation and the further consumption of the degraded ureide products, as indicated by the low level of allantoin in the WT and their enhancement in the old leaves of the low nitrate fed *Ataln* plants (Fig. 4B). Importantly, the negligible allantoin levels evident in *Atxdh1* leaves, which are at a much lower level than in WT leaves (Fig. 4B), was shown recently by others (Brychkova *et al.*, 2008; Watanabe *et al.*, 2014) and can be explained by a slight XDH activity resulting from AtXDH2 or another yet unknown source. Importantly, the levels of allantoin accumulated in both young and old *Ataln* leaves were much higher than the level of xanthine accumulated in *Atxdh1* leaves, indicating that the range between the accumulation of xanthine and allantoin, may be an indication for the rate of purine catabolism in WT plants.

Interestingly, while a large difference between older and younger leaves is evident for xanthine accumulation, the ureide accumulation was much less differential (Fig. 4, Supplementary Fig. S3). This indicates that while the initial purine breakdown takes place in the older leaves, the allantoin is readily exported and the subsequent catabolic steps take place in all leaves. This scenario is consistent with the observation of elevated UPS transcript levels in WT older leaves (Fig. 3).

### The expression of the purine degradation network is up regulated by nitrogen deficiency and down regulated by high nitrate application

During natural senescence and dark induced senescence an orchestrated regulation of transcripts related to purine catabolism was observed that includes upregulation of upstream purine catabolism transcripts such as *Atxdh1* and *AtUOX* with parallel downregulation of the downstream transcripts *AtALN* and *AtAAH* (Brychkova *et al.*, 2008). We wished to elucidate how the purine catabolism gene network is orchestrated either by N-deficiency induced senescence and/or by sufficient N that prevents senescence in purine mutant leaves. Therefore, the transcripts of the purine-degrading enzymes were analyzed in plants supplemented with low and high nitrate as the only N source. The low nitrate as compared to high nitrate treatment resulted in the increase in transcript expression of purine catabolism genes in WT plants. Whereas *AtAMPD*, *AtAAH* and *AtALN* were increased in old and young leaves, *Atxdh1 AtUOX* were found to increase only in old leaves (Fig. 5). Interestingly, with the exception of the *Atxdh1* transcript, *Atxdh1* mutant showed a similar tendency towards the expression pattern of the purine degradation gene network as the WT (Fig. 5). Importantly, *Ataln* and *Ataah* mutants also exhibited a similar general expression tendency as WT and *Atxdh1* mutant (Supplementary Fig. S4, Fig. 5). Thus, we can conclude that the purine degradation transcript network is generally up regulated by nitrogen deficiency and down regulated by high nitrate application, indicating that the regulation in senescent leaves is more complex.

**Figure 5.**
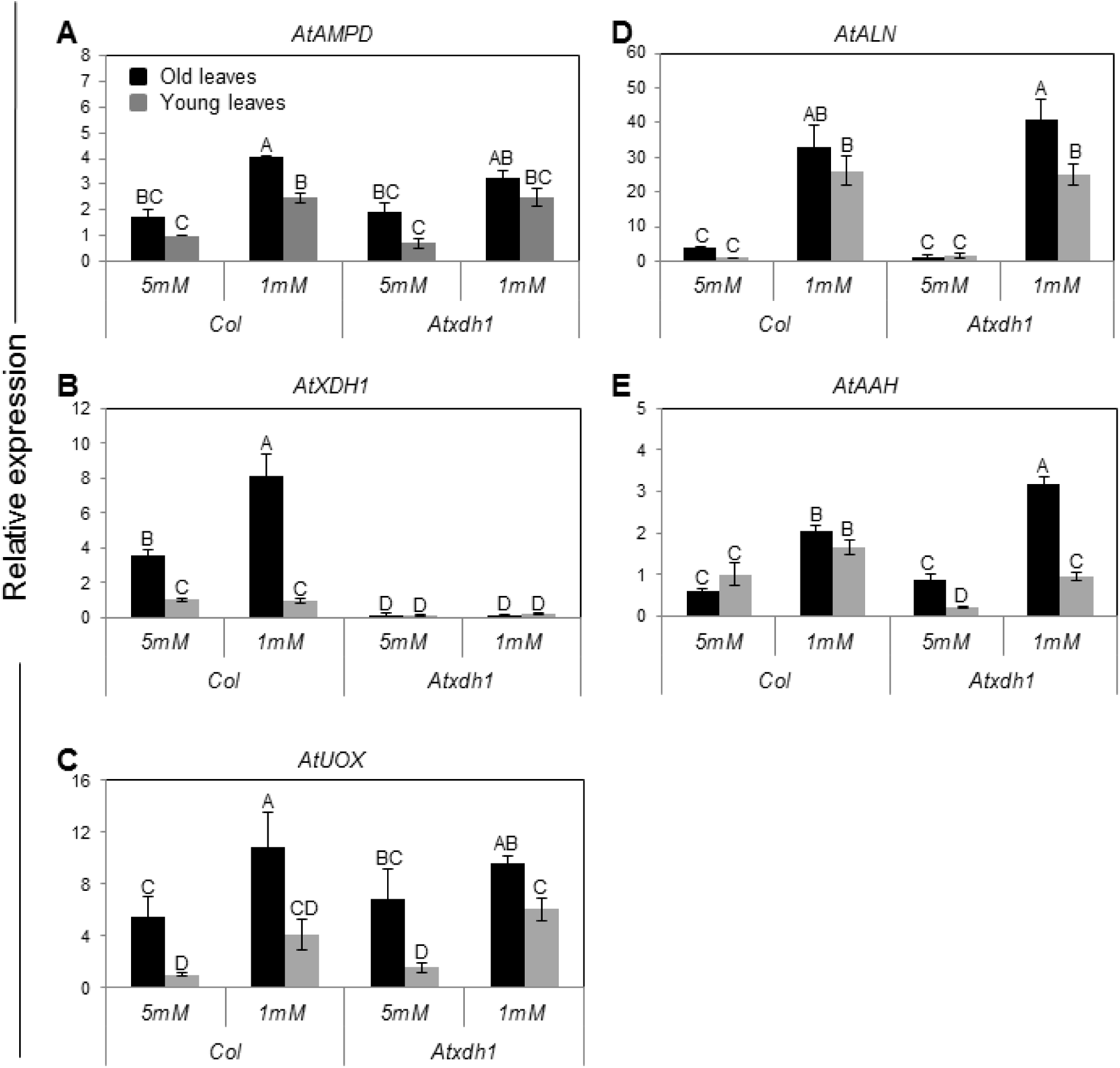
Transcript expression of the purine catabolism genes in young and old leaves of WT and *Atxdh1* plants supplemented with 1 or 5mM nitrate as the only N source. Adenosine 5’-monophosphate deaminase (*AtAMPD*) (A), xanthine dehydrogenase (*AtXDH*) (B), urate oxidase (*AtUOX*) (C), allantoinase (*AtALN*) (D) and allantoate amidinohydrolase (*AtAAH*) (E) were presented. Quantitative analysis of transcripts by real-time RT-PCR was performed using WT (Col) and *Atxdh1* (SALK_148366. GABI_049004) 25 days old plants grown on nitrogen deficient soil. The expression of each treated line was compared with the young leaves of WT in 5mM nitrate treatment after normalization to *EF-1α* gene product (At5g60390). The data represent the mean obtained from three independent experiments. Values denoted by different letters are significantly different (Tukey-Kramer HSD test, P < 0.05).

### Low nitrate supplementation confers enhanced XDH activity and AAH protein in leaves

To examine the levels of protein expression we chose *XDH1* and *AAH,* as representative proteins of the upstream and downstream components of the purine catabolism, to be estimated by activity gels and immunodetection, respectively (Fig. 6, Supplementary Fig. S5 A). The results indicated elevated levels of XDH activity in low nitrate in both old and young leaves (Fig. 6A). NADH dependent superoxide generating activity of XDH, confirmed this result (Supplementary Fig. S5 B).

**Figure 6.**
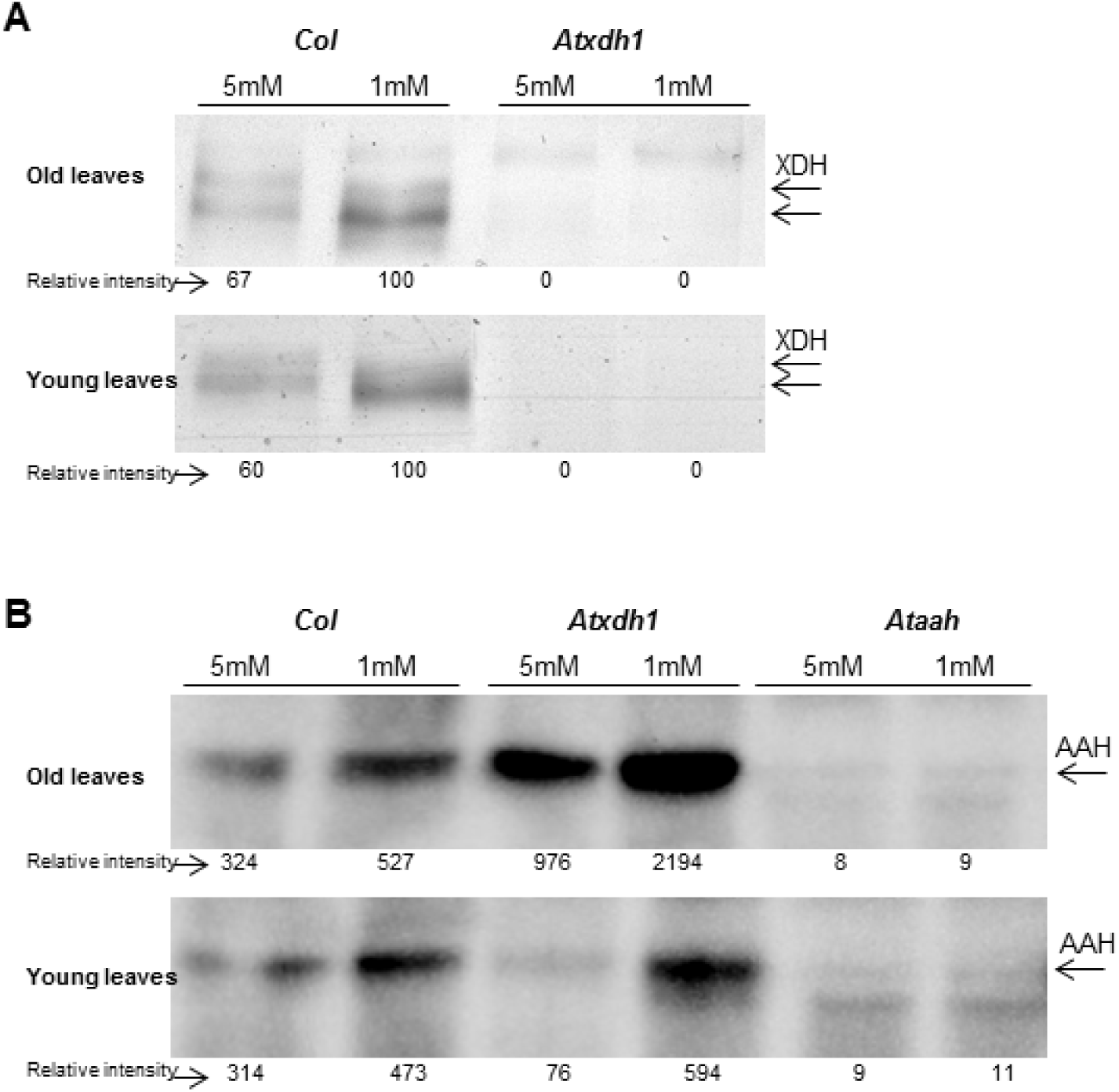
The activity of xanthine dehydrogenase (XDH) (A) and immunoblot analysis of allantoate amidohydrolase (AAH) (B). Protein extracted from old and young leaves of WT (Col) and *Atxdh1* mutant (SALK_148366, GABI_049004) grown on 1 mM or 5 mM nitrate as the only N source. The general activity of XDH in native-SDS PAGE gel was detected by using PMS as the electron-carrier intermediate and MTT as the electron acceptor. AAH protein level was analyzed by immunoblotting employing specific antiserum against AAH. Protein extracted from *Ataah* (SALK_112631) mutant leaves was used as the negative control. For XDH activity 50 and 100 μg crude protein extracts was loaded per each lane for the old and young leaves. For AAH analyses equal amount of 100 μg crude protein extracts were loaded per each lane for old and young leaves respectively. The data represents one of three independent experiments with similar results.

AAH protein expression was evaluated by immunoblotting after native PAGE, employing highly specific antisera (gifted by Claus-Peter Witte). The AAH protein was detected as one band in the WT, verified by being absent in *Ataah* (Fig. 6B). Interestingly, analyses of AAH levels in WT and in the *Atxdh1* mutant revealed that the AAH level increased in old and young leaves with decreasing nitrate supplementation (Fig. 6B). These results indicate that XDH1 and AAH protein and activity expression are generally in agreement with the results of the transcript network expression and indicates generation of ureides and their degradation for further deployment of nitrogen during its deficiency (Fig. 6A, B).

### Nitrate reductase activity in 18 and 25 days old *Atxdh1* and WT plants fed with high and low nitrate levels

Nitrate reductase (NR) catalyzes the first step of nitrate assimilation toward the biosynthesis of NH3, by generating the intermediate nitrite (Campbell, 1988; Kaiser and Huber, 1994; Sivasankar and Oaks, 1996). To examine the influence of purine catabolism potential, NR activity was estimated in young and old leaves of 18 days old WT and *Atxdh1* mutant plants supplemented with high and low nitrate as the only N source (Fig. 7A). In general NR activity was significantly higher in younger leaves than in older leaves. These results are consistent with the finding that primary N-assimilating enzymes decrease with aging, as shown before (Masclaux et al. 2000). In addition, the level of NR activity was always more elevated in the mutant line. Lower nitrate levels were detected in the young as compared to the old leaves of 25 and 18 days old plants (Fig. 2C and Fig. 7 B, respectively) and in the *Atxdh1* mutant compared to the WT in the 18 days old plants (Fig. 7B) indicating a higher nitrate assimilation by NR (Fig. 7A). The higher NR activity in the mutant younger leaves, followed by lower nitrate, may indicate compensation for the lack of remobilized purine-dependent nitrogen source from the older to younger leaves, especially in the nitrate starved *Atxdh1* mutant plants.

**Figure 7.**
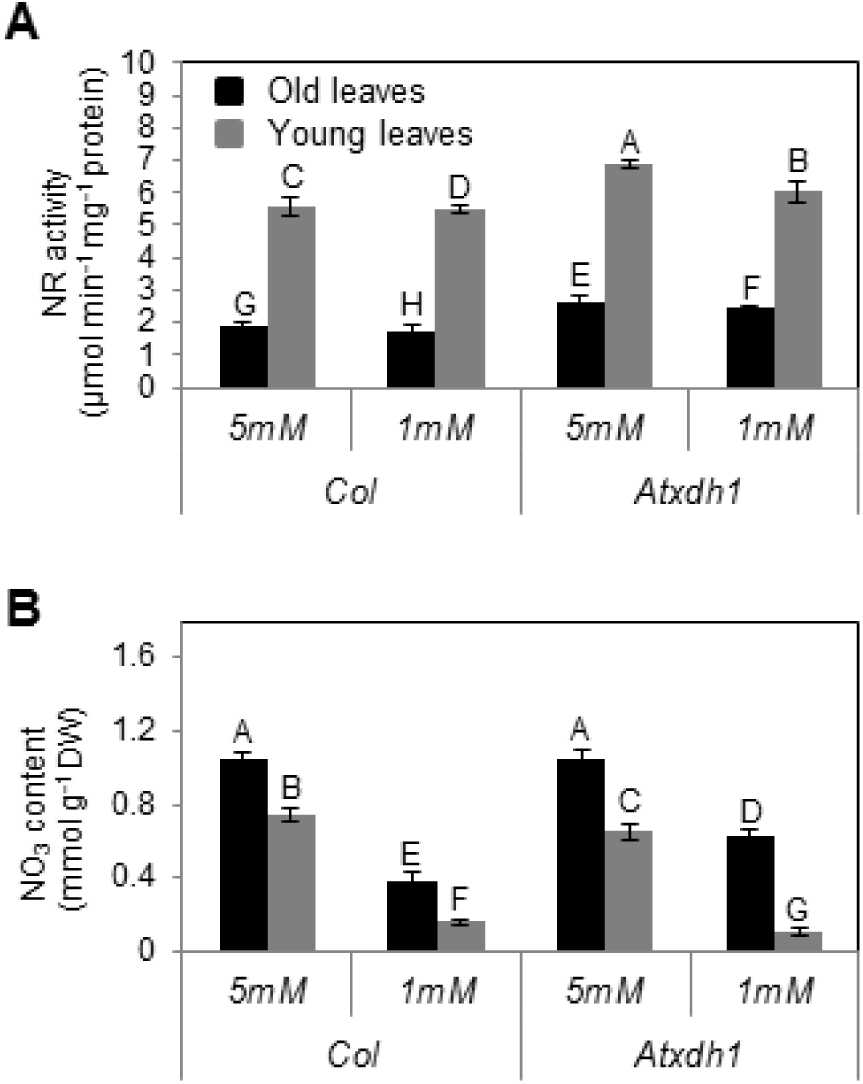
Nitrate reductase activity (A) and nitrate content (B) in old and young leaves of WT (Col) and *Atxdh1* mutant as affected by nitrate level supplemented to the growth medium. Eighteen days old plants grew in nitrogen deficient soil supplemented with one-half-Hoagland nutrient solution containing 1 or 5mM NaNO_3_ as the only N source. The data represent the mean obtained from three independent experiments. Values denoted by different letters are significantly different (Tukey-Kramer HSD test, P < 0.05). *Atxdh1* are GABI_049D04 and SALK_148366 T-DNA mutants

## DISCUSSION

In tropical legumes such as soybean, cowpea and common bean, symbiotic N fixation in plant nodules leads to the incorporation of the fixed N into purine nucleotides to be converted into ureides, which are used as major nodule (root) to shoot nitrogen transport compounds (Schubert, 1986; Amarante *et al.*, 2006). In other plants, leaf senescence has been shown to be accompanied by a decrease in nucleic acids content (Masclaux *et al.*, 2000). Hence, it is reasonable to consider that relatively N enriched ureides (1:1 C/N ratio) could serve as possible endogenous conduits for remobilized nitrogen during plant growth and development in non-legumes (Have *et al.*, 2016). In support of this notion is the observation that ureides accumulate in several non-legume shrubs and trees, likely for storage and translocation of nitrogen (Schmidt and Stewart, 1998).

The presence of all components necessary for purine catabolism and its recycling and remobilization are well established in non-legumes plants (Todd and Polacco, 2006; Todd *et al.*, 2006; Zrenner *et al.*, 2006; Werner and Witte, 2011). However, the physiological role of purine degradation, how it contributes to N remobilization and overall N nutrition and the conditions under which it becomes a critical limiting step in organ survival are not known.

By analyzing the central mutations of the purine degradation pathway under nitrogen-limiting conditions, we show the critical role of the purine degraded product in providing nitrogen. Based on this work, one general physiological role of purine degradation in plants is to optimize the use of N resources by recycling purines from N source tissues such as older leaves for remobilization to growing organs and tissues as N-sinks.

### The integration of nitrogen starvation and impairment in purine catabolism results in early senescence symptoms in old leaves of Arabidopsis plants

Leaf senescence hallmarks - decrease in chlorophyll level, soluble protein content, organic nitrogen content and enhancement of senescence molecular markers were evident in the Arabidopsis mutant plants *Atxdh1*, *Ataln* and *Ataah* supplemented with 1mM nitrate as the only N source (Fig. 1, 2, Supplementary Fig. S1). Leaf senescence has an important role in nitrogen metabolite management, to remobilize important degraded N containing components for the re-assimilation of nitrogen resulting from chloroplast degradation, hydrolysis of stromal proteins and other degraded organelles and cell components (Masclaux *et al.*, 2000; Hörtensteiner and Feller, 2002; Eckhardt *et al.*, 2004; Fischer, 2007; Liu *et al.*, 2008). A similar metabolic strategy was suggested to take place in senescence that resulted from nitrogen limitation (Aerts, 1990). Importantly, old leaves of WT plants that undergo nitrate starvation do not show senescence symptoms whereas the various independent mutants; *Atxdh1, Ataln* and *Ataah,* impaired in the purine catabolism pathway did (Fig. 1, 2, Supplementary Fig. S1). Senescence symptoms and elevated expression of the senescence marker genes *SAG12*, *SGN1* and *ACD2* were noted in old leaves of the purine pathway mutants grown with low nitrate (Fig. 1, Supplementary Fig. S1). These leaves also exhibited significantly higher expression of *GLN1.1*, *GDH1*, *GDH2* transcripts as compared to WT (Supplementary Fig. S6). Enhanced expression of the latter transcripts was shown previously to be associated with protein degradation necessary to remobilize nitrogen from senescent leaves in tobacco (Masclaux *et al.*, 2000; Pageau *et al.*, 2006). Considering the absence of all these described senescence symptoms in old leaves of nitrate starved WT one must conclude that the absence of purine catabolism forces the plant to compensate for nitrogen shortage by activating chloroplast proteins degradation in the old leaves (Matile *et al.*, 1996; Suzuki and Shioi, 1999; Pruzinska’ *et al.*, 2005). Hence, a robust catabolic pathway is necessary to prevent premature senescence.

### Allantoin, a degraded N enriched purine metabolites is remobilized from the old to the young growing leaves

The lack of senescence symptoms in the old leaves of nitrate starved WT plants, whereas *Atxdh1*, *Ataln* and *Ataah* mutants exhibited senescence symptoms that could be prevented by enhanced nitrate supplementation (Fig. 1 and Supplementary Fig. S1), indicate the essentiality of a normal active purine catabolism pathway for an efficient nitrogen metabolism. Importantly, while the decrease in soluble protein and organic N in the old leaves of nitrate starved *Atxdh1* mutant could be avoided by supplementation of 5 mM nitrate, a significantly lower nucleic acid level, expressed as total RNA, was detected in old leaves of the nitrate starved WT, whereas no senescence symptoms were noticed (Fig. 2). These results support a notion of remobilization of the degraded nucleic base purine metabolites such as ureides from the old to young leaves. This notion was additionally supported by allantoin infiltration by injection into the old leaves of WT and *Atxdh1* mutant, where a significant enhancement of allantoin levels was evident not only in the injected leaves, but in the middle and the young leaves of WT as well as *Atxdh1* mutant as compared to the control (H_2_O) infiltrated plants. Importantly, the accumulated allantoin was significantly higher in the youngest than in the middle leaves (Supplementary Fig. S7). This scenario is further supported by the significant enhanced expression of *AtUPS1*, *AtUPS2* and *AtUPS5* transporters in old WT leaves of nitrate starved plants compared to plants supplemented with high nitrate or to *Atxdh1* mutant plants (Fig. 3). The up-regulation of the nitrogen-related transporters was previously suggested as an indication that the transporters participate in the remobilization of the related nitrogen forms from the senescing tissues (Kojima *et al.*, 2007). Hence, enhanced expression of the ureide transporters is indicative of ureide remobilization from the old leaves to the young growing leaves of nitrate starved plants. Furthermore, the enhanced transcript levels of the upstream and downstream purine catabolism genes as well as their protein levels were preferentially detected in old leaves of nitrate starved plants (Fig. 5, 6 and Supplementary Fig. S4). Such orchestration of purine catabolism should normally result in full degradation of purine metabolites and indeed, xanthine and allantoin do not accumulate in WT leaves (Fig. 4), whereas they do accumulate in their related mutants. Higher accumulation of xanthine was evident in the old as compared to the young leaves of *Atxdh1* mutants and significantly higher allantoin was noticed in the young leaves of *Ataln* mutant. This indicates that xanthine is mostly degraded in the old leaves and the majority of the generated allantoin is remobilized to the young leaves (Fig. 4). The results support the notion of ureide remobilization taking place from the old leaves to the young growing leaves.

### Higher protein degradation during the period of 18 to 25 day after germination is the cause for pre-mature senescence symptoms in the older leaves of nitrate starved *Atxdh1* plants

N remobilisation was shown to start earlier and with increased rate of the remobilized N when plants grow with low as compared to high nitrogen (Ta and Weiland, 1992; Uhart and Andrade, 1995). Emerging and growing organs, such as young leaves, are a potential sink to trigger N remobilisation from older plant parts, that includes N remobilisation from leaf to leaf during the vegetative phase (Wendler *et al.*, 1995; Masclaux-Daubresse *et al.*, 2008). Proteins in the mature leaves are potential N storage to be degraded and remobilized to the young growing leaves (Hensel, 1993; Masclaux *et al.*, 2000; Hörtensteiner and Feller, 2002; Fischer, 2007; Liu *et al.*, 2008). Amongst these, are chloroplastic proteins such as Rubisco, which represent 50% of the total proteins in mature leaves of C3 plants (Staswick, 1997). Indeed, Rubisco large subunit and D1 protein (encoded by *psbA* transcript), a component of the reaction center of PSII (Keren *et al.*, 1997), both chloroplast releted proteins, were decreased in old leaves of nitrogen limited *Atxdh1* mutant, whereas, the level of the autophagy-related protein ATG8a was enhanced, indicating enhanced remobilization of the degraded protein products (Supplementary Fig. S8). This is consistent with the observation of the 33% or even more than the 45% reduction of the soluble protein content in these leaves as compared to the older leaves of the WT starved plant or the old leaves of the elevated nitrate fed plant, respectively (Fig. 2E).

Importantly, no difference in chlorophyll content was evident, and no yellowish of leaves was noticed in the old leaves of the nitrate starved *Atxdh1* plants at the age of 18 days (Supplementary Fig. S9A and B). Furthermore, at this growth stage the mutation in *Atxdh1*, or the level of the supplemented nitrate had no effect on the level of the soluble proteins in the old leaves (Supplementary Fig. S9 C). Considering that the level of the soluble proteins decreased in the older leaves and increased in the younger leaves of the 25 as compared to the 18 days old plants (compare Supplementary Fig. S9C to Fig. 2E), our results indicate a higher soluble protein degradation and remobilization rate from the older to the younger leaves of the nitrate starved *Atxdh1* than WT plants. Further considering that the nitrogen-to-protein conversion factor ranged in plant leaves from 6.25 to 4.43 (Yeoh and Wee, 1994) the decrease of 0.13 mg protein g^-1^ FW indicates the remobilization of 20.8 to 29.3 μg more N originating from degraded soluble protein, in N starved mutant older leaves as compared to WT (Fig. 2E). Importantly, the 0.37 or 0.78 μmol g^-1^ FW xanthine or allantoin (Fig. 4), contains 20.71 or 43.65 μg N g^-1^ FW (see calculation in Supplemental Table S2) accumulated in the old leaves of nitrate starved *Atxdh1* and *Ataln* mutants respectively, that likely represents the rate of purine catabolism and remobilization of the resulting N from older to younger growing WT leaves. Given so, the results clearly demonstrate that the absence of the purine degraded N remobilized from the older leaves is the cause for the senescence symptoms, a result of higher soluble protein degradation, several days before bolting, in older leaves of nitrate starved *Atxdh1* plants.

### The level of the applied nitrate negatively regulates purine degradation in Arabidopsis leaves

The application of nitrate was shown to downregulate *AtALN* and *AtAAH* transcripts in two nitrogen starved Arabidopsis plants (Werner *et al.*, 2008) and inhibit nodule formation as well as the fixation of atmospheric N_2_ in legume plants (Murray et al., 2016 and references therein) indicating that nitrate supplementation is associated with purine metabolism and negatively affects purine metabolite use in legumes and non-legume plants. This was clearly demonstrated by the lower accumulated xanthine and allantoin in leaves of 5 mM nitrate supplemented *Atxdh1* and *Ataln* than in the 1 mM nitrate fed mutants, respectively (Fig. 4). Nitrate supplementation is shown here to negatively regulate purine catabolism at both the transcript (Fig. 3, 5, 7) and protein (Fig. 6 and Supplementary Fig. S5) expression levels. However, another level of posttranslational modification was shown before to exist in ryegrass. In that case, the lower XDH activity as well as lower allantoin and allantoate levels in leaves of nitrate supplied annual ryegrass as compared to ammonium supplied plants (Sagi *et al.*, 1998), were attributed to the preferred allocation of the molybdenum cofactor (Moco). The catalytic center of NR and XDH1, plant molybdoenzymes, requires Moco. Preferential allocation of Moco to NR supports nitrate assimilation in the presence of high nitrate over AtXDH1 activity (Sagi *et al.*, 1997, 1998; Sagi and Lips, 1998). While not examined here directly, the enhanced AtXDH1 activity and decreased NR activity in old leaves and vice versa in young leaves, as well as the enhanced AtXDH1 activity and decreased NR activity in nitrate starved plants and vice versa in leaves of high nitrate supplied plants, supports this notion (Fig. 6, 7, Supplementary Fig. S5). This study uncovers further transcriptional and post-transcriptional levels of control.

High nitrate application almost fully abrogated the senescence symptoms evident in the old leaves of nitrate starved mutants, by enhancing organic nitrogen level and soluble protein content in these leaves (Fig. 1, 2 and Supplementary Fig. S1). This is attributed to NR activity that was significantly higher in mutant than in WT plants when detected in leaves of 18 day old plants (Fig. 7), before the appearance of senescence symptoms (Supplementary Fig. S9). The lower nitrate (Fig. 2, 7B) and higher NR activity in *Atxdh1* mutant than in WT leaves when both plants were supplemented with high nitrate (Fig. 7) suggest that under high nitrate availability the shortage of purine originated nitrogen in *Atxdh1* mutant plant is compensated by higher nitrate assimilation.

## MATERIALS AND METHODS

### Plant Material Growth Conditions

*Arabidopsis thaliana* (L.) Heynh wild-type and mutants used in the current study were of the Col-0 background. The following homozygous T-DNA inserted mutants were employed: *Atxdh1*(GABI_049D04, SALK_148366, accession No. At4g34890) described previously by us (Yesbergenova et al. 2005, Brychkova et al., 2008); *Ataln* (SALK_ 013427, SALK_146783, accession No. At4g04955) described before (Todd and Polacco 2006; Watanabe et al., 2014); *Ataah* (SALK_112631; Todd & Polacco 2006, accession No. At4g20070).

Seeds were surface-sterilized in 80% alcohol for 2 min, washed three times in sterile water and sown on one-half strength Murashige and Skoog (MS 1/2) agar plates (Murashige and Skoog, 1962). The plates were placed at 4°C for 3 days to synchronize germination, and then were transferred to a controlled growth room at 22°C, 14/10 h light/dark photoperiod and light intensity of 150 μE m^−2^ s^−1^. Six-day-old seedlings were transferred each to a 0.128 L pot containing a 1:1 mixture of perlite and nutrient-free soil (Sun Gro Horticulture Canada; http://www.sungro.com/). Plants were irrigated twice a week with a 0.5 Hoagland solution (Hoagland and Arnon, 1950) modified to contain 1 or 5 mM NaNO_3_, as the only N source, where the sodium level was balanced to contain 5 mM sodium in all the treatments, by the supplementation of NaCl. Salinization was avoided by irrigation performed to leach out 50% of the irrigated nutrient solution. The leaves of plants at 18 or 25 days after germination (The latter just before bolting) were harvested, snap-frozen in liquid nitrogen and stored at −80°C for further use. The first 4 rosette leaves from the bottom were designated as old leaves and the upper-most 4 from top as young leaves.

### Determination of chlorophyll and anthocyanin

For chlorophyll determination, 4 leaf discs were sampled from old and young rosette leaves of WT, *Atxdh1*, *Ataln*, *Ataah* mutants grown under the low and high nitrate conditions. The leaf discs (0.7mm diameter) were immersed in 90% EtOH and incubated at 4°C for 2 days in the dark. Absorbance of the extracted chlorophyll was measured at 652 nm and total chlorophyll was estimated (Ritchie, 2006). To assess the response to external xanthine and allantoin, 7mm leaf discs were sampled from rosette leaves of WT and *Atxdh1* mutant plants and put in Petri dishes, on filter paper soaked with a solution containing water (mock) and 1 mM xanthine or allantoin, for two days in permanent light. Thereafter the discs were washed and the total anthocyanin was measured as described in Laby et al., 2000. In addition, the green area of the leaf disk was estimated by employing Digimizer 3.2.1.0 (http://www.digimizer.com) and presented as the ratio of the green part to total area of the leaf disc, as the chlorophyll damage indicator.

### Metabolites Analysis

Samples (100 mg) were grounded in 25 mM pH 7.5 K_2_PO_4_/KH_2_PO_4_ buffer (1:4 w/v) using a chilled mortar and pestle (Brychkova *et al.*, 2008; Lescano *et al.*, 2016). The resulting homogenates were transferred to 1.5 ml micro-centrifuge tubes, centrifuged at 15000 *g* for 20 min at 4°C, and the supernatant was used for analyses. Quantification of the ureides, allantoin and allantoate was performed using the differential conversion of ureide compounds to glyoxylate and colorimetric detection described by Vogels and Van Der Drift, (1970) and employed by others (Todd *et al.*, 2006; Brychkova *et al.*, 2008; Werner *et al.*, 2008, 2013; Watanabe *et al.*, 2014; Lescano *et al.*, 2016; Takagi *et al.*, 2016). Xanthine was detected using the xanthine oxidase assay as previously described (Brychkova et al., 2008a). Ammonium was detected by Nessler method (Molins-Legua *et al.*, 2006). Nitrate content was analyzed according to Cataldo et al. (1975). Total N in the dried tissues was measured by an elemental analyzer (Thermo Scientific™ FLASH 2000 CHNS/O Analyzers).

### Protein Extraction and Fractionation

Whole protein from Arabidopsis rosette leaves was extracted as described by Sagi et al., (1998). Concentrations of total soluble protein in the resulting supernatant were determined according to (Bradford, 1976). Native-SDS PAGE was carried out as previously described (Sagi and Fluhr, 2001; Srivastava *et al.*, 2017). Samples containing the extracted proteins were incubated on ice for 30 min in sample buffer containing 47 mM Tris-HCl (pH 7.5), 2% (w/v) SDS, 7.5% (v/v) glycerol, 40 mM 1,4-dithio-DL-threitol (DTT) as the thiol-reducing agent, and 0.002% (w/v) bromophenol blue. The incubated samples were centrifuged at 15,000xg for 3 min before loading the supernatant and subsequently resolved in 7.5% (w/v) SDS-polyacrylamide separating gel and 4% (w/v) stacking gels. Native-SDS PAGE was carried out using 1.5 mm thick slabs loaded with 50 μg of old leaf or 100 μg of young leaf proteins unless mentioned otherwise.

### XDH In-gel activity and nitrate reductase kinetic activity

Regeneration of the active proteins after denaturing PAGE was carried out by removal of the SDS by shaking the gel for 1 h in 10mM Tris-HCl buffer (pH 7.8) solution (65 ml buffer per ml of gel) containing 2mM EDTA and 1.0% (w/v) Triton X-100 (Sagi and Fluhr, 2001; Srivastava *et al.*, 2017). Following the regeneration process, the gels were assayed for normal in-gel XDH activities using 0.1mM PMS, 1mM MTT and addition of 0.5mM xanthine mixed with 1mM hypoxanthine in 0.1mM Tris-HCl buffer (pH 8.5), at 25 °C under dark conditions. To detect superoxide generation activity of XDH, PMS was omitted and the mix of xanthine with hypoxanthine or 0.25 mM NADH as specific substrates were employed (Yesbergenova *et al.*, 2005). The quantity of the resulting formazan was directly proportional to enzyme activity during a given incubation time, in the presence of excess substrate and tetrazolium salt (Rothe, 1974; Srivastava *et al.*, 2017).

For nitrate reductase activity the samples were extracted in a buffer containing 3 mM EDTA, 3.6 mM dithiothreitol (DTT), 0.25 M Tris-HCl (pH 8.48), 3 mg L-Cys, 3 mM NaMoO4 and protease inhibitors including aprotenin (10μg ml^-1^) and pepstatin (10μg ml^-1^) and the activity was detected as previously described (Sagi et al., 1997).

### Western Immunoblotting

Protein crude extract samples (20–50*μ*g) extracted as described by Sagi et al., (1998), were subjected to Native-PAGE for detecting AAH and SDS-PAGE electrophoresis for the other proteins. The fractionated proteins were transferred onto polyvinylidene difluoride membranes (Immun-Blot membranes, Bio-Rad). The membrane was probed first with the following primary antibodies: Anti-AAH (a gift from ClausPeter Witte, http://www.ipe.uni-hannover.de) at 1:500 dilution ratio, antibody specific to D1 protein [a component of the reaction center of PSII] (Agrisera, http://www.agrisera.com) at 1:10,000 ratio, specific antibodies to autophagy-related protein 8a (ATG8a) (Abcam, http://www.abcam.com) at 1:1,000 ratio and an antibody recognizing large subunits of Rubisco (LSU) (a gift from Michal Shapira (http://in.bgu.ac.il/en/natural_science/LifeSciences/Pages/staff/Michal_Shapira.aspx) at a dilution ration of 1:3,000. Thereafter, the proteins under went binding with secondary antibodies diluted 5000-fold in PBS (anti-rabbit IgG; Sigma-Aldrich). Protein bands were visualized by staining with Clarity Western ECL Substrate (Bio-Rad, USA) and quantified by Image lab (Version 5.2, Bio-Rad, USA).

### Quantitative RT-PCR

Total RNA was extracted from plants using the Aurum Total RNA kit according to the manufacturer’s instructions (Bio-Rad). First-strand cDNA was synthesized in a 10-μl volume containing 350 ng of plant total RNA that was reverse transcribed employing an iScript cDNA Synthesis Kit (Bio-Rad) according to the manufacturer’s instructions. The generated cDNA was diluted 10 times, and quantitative analysis of transcripts was performed employing the set of primers presented in Supplemental Table S1 designed to overlap exon junctions as previously described (Brychkova et al., 2007).

### Statistical analysis

All results are presented as means and standard errors of means. The data for total N, total organic nitrogen, nitrate, and ammonium represent the mean obtained from six independent experiments. Metabolite, protein content and transcripts measurements represent mean obtained through at least three independent experiments. Each treatment was evaluated using ANOVA (JMP 8.0 software, http://www.jmp.com). Comparisons among three or more groups were made using one-way analysis of variance with Tukey’s multiple comparison tests.

## SUPPLEMENTARY DATA

**Supplemental Figure S1**. The effect of nitrate level supplemented to the growth medium of wild-type (WT) and mutant plants impaired in the purine catabolic pathway.

**Supplemental Figure S2**. Effects of exogenous application of xanthine and allantoin on leaf disc appearance.

**Supplemental Figure S3**. The levels of the xanthine content in old and young leaves of the *Atxdh1*, *Ataln* and *Ataah* mutant and WT plants grown in nitrogen deficient soil supplemented with 1mM NaNO_3_.

**Supplemental Figure S4.** Transcript expression levels of purine catabolism transcripts in young and old leaves of WT (Col) and purine catabolism impaired plants supplemented with 1 mM or 5 mM nitrate as the only N source.

**Supplemental Figure S5**. Xanthine and NADH depended superoxide-generating activities of XDH in old and young WT and *Atxdh1* mutant leaves of plants grown with 1 mM or 5 mM nitrate as the only N source.

**Supplemental Figure S6**. The relative expression of N assimilation senescence related transcripts in old and young leaves of WT and *Atxdh1* mutant plants.

**Supplemental Figure S7**. The effect of allantoin infiltration to the oldest leaves on its level in old, milled and young leaves of WT and *Atxdh1* mutant plants.

**Supplemental Figure S8**. Immunoblot analysis of large subunit of Rubisco (LSU), component of the reaction center of PSII D1 and autophagy protein ATG8a.

**Supplemental Figure S9**. Leaf appearance, total chlorophyll content and soluble protein content of the old and young leaves of 18 days old WT and *Atxdh1* mutant plants.

**Supplemental Tables**

**Supplemental Table S1**. Gene-specific primer sequences used for the expression analyses

**Supplemental Table S2**. The calculation of nitrogen content in xanthine and allantoin accumulated in the old leaves of *Atxdh1* and *Ataln* mutants respectively.

## List of author contributions

A.S. participated in designing the research plans and performed the experiments and analyses; S.S participated in XDH in gel assay; A.K. participated in nitrate and ammonium detection; A.B. participated in qRT-PCR. R.F. read and commented on the manuscript. M.S. conceived the original idea, designed the research plan, and supervised the research work. The manuscript was jointly written by A.S. and M.S.

## ACKNOWLEDGEMENTS

This research was supported by the Israel Center of Research Excellence (ICORE) ‘Plant Adaptation’ (ISF Grant no. 757/12). Dr Sudhakar Srivastava is the recipient of a postdoctoral fellowship from the Jacob Blaustein Center for Scientific Cooperation. We thank Dr. Tali Bruner for editing the manuscript.

## Supplemental Figures

**Supplemental Figure S1**. The effect of nitrate level supplemented to the growth medium of wild-type (WT) and mutant plants impaired in the purine catabolic pathway. Leaf appearance (A) and total chlorophyll content (B) of the old leaves of WT (Col), *Atxdh1* (GABI_049D04), *Ataln* (SALK_146783) and *Ataahl* (SALK_112631) mutant plants. Plants were grown in nitrogen deficient soil supplemented with 1 or 5 mM NaNO_3_ as the only N source. Relative transcript expression levels of suppressor of overexpression of Cys protease senescence-associated gene 12 [*SAG12*, (At5G45890)] (C), stay-green protein 1 [*SGN1* (At4G22920)] (D), in the old leaves of WT and mutants is presented. Transcript levels were compared to the expression in 5mM NaNO_3_ treated WT old leaves, after normalization to Elongation factor alfa (*EFα*) gene (At5G60390) and presented as relative expression. The data represents the mean obtained from three independent experiments. Values denoted by different letters are significantly different (Tukey-Kramer HSD test, P < 0.05).

**Supplemental Figure S2**. Effects of exogenous application of xanthine and allantoin on leaf disc appearance. Appearance of adaxial (left insert) and abaxial (right insert) surface is presented (A). Leaf discs removed from 6^th^ to 10^th^ rosette leaves [counted from the bottom (without senescence symptoms)] of WT and *Atxdh1* (SALK_148366) mutant plants were treated with water (mock) and 1 mM xanthine or allantoin under permanent light (for 48 h). The green area ratio of adaxial (B) and abaxial (C) side was estimated by Digimizer 3.2.1.0 tool (http://www.digimizer.com). Discs were washed and the total anthocyanin content (D) was determined. The presented data is one of three independent experiments with similar results. Values denoted by different letters are significantly different (Tukey-Kramer HSD test, N=3, P < 0.05).

**Supplemental Figure S3**. The levels of the xanthine content in old and young leaves of the *Atxdh1*, *Ataln* and *Ataah* mutant and WT plants grown in nitrogen deficient soil supplemented with 1mM NaNO_3_. The data represent the mean obtained from three experiments. The values denoted with different letters are significantly different according to the Tukey-Kramer HSD test; p< 0.05. Different upper case letters indicate differences between mutants and WT plants. The following T-DNA mutants were emplpyed: *Atxdh1,* SALK_148366 and GABI_049004; *Ataln*, SALK_013427 and SALK_146783; *Ataah*, SALK_112631.

**Supplemental Figure S4**. Transcript expression levels of purine catabolism transcripts in young and old leaves of WT (Col) and purine catabolism impaired plants supplemented with 1 mM or 5 mM nitrate as the only N source. The expression of Adenosine 5’-monophosphate deaminase (*AtAMPD*) (A), xanthine dehydrogenase (*AtXDH*) (B), urate oxidase (*AtUOX*) (C), allantoinase (*AtALN*) (D) and allantoate amidinohydrolase (*AtAAH*) (E) were presented. Quantitative analysis of transcripts by realtime RT-PCR was performed using 25-days-old plants. The expression of each treated line was compared with the young leaves Col grown with 5 mM nitrate after normalization to *EF-1α* gene product (At5g60390). The data represent the mean obtained from three different experiments. Values denoted by different letters are significantly different (Tukey-Kramer HSD test, P < 0.05). The following T-DNA mutants were employed: *Atxdh1*, GABI_049D04; *Ataln*, SALK_146783; *Ataah*, SALK_112631.

**Supplemental Figure S5**. Xanthine (A) and NADH depended (B) superoxide-generating activities of XDH in old and young WT and *Atxdh1* mutant leaves of plants grown with 1 mM or 5 mM nitrate as the only N source. Xanthine dependent XDH assay contained MTT and hypoxanthine/xanthine, NADH depended XDH assay contained NADH and MTT. The data represents one of 3 independent experiment with similar results. The lanes of old and young leaves of the activity gels contained 50 and 100 μg of crude protein extract, respectively. SALK_148366 was employed as the T-DNA mutant of *Atxdh1*.

**Supplemental Figure S**.6 The relative expression of N assimilation senescence related transcripts in old (A) and young leaves (B) of WT (Col) and *Atxdh1*(SALK_148366) mutant plants. The 25-days-old plants grew in N-deficient soil containing 1 or 5mM NaNO3 as the only N source. The relative expression of the following transcripts were evaluated: *Gln1;1*, *Gln1;2*, *Gln1;3*, *Gln1;4*, *Gln1;5*, *GDH1*, *GDH2* and *GDH3*. The expressed transcript levels were compared to *Gln1;5* and *GDH3* expression, respectively, in young leaves of Col grown in 5mM NaNO3, after normalization to the Elongation factor alfa (*EFα*) gene (At5g60390.1) and presented as relative expression for *Gln* and *GDH*, respectively. The data represent the mean obtained from three independent experiments. Values denoted by different letters are significantly different (Tukey-Kramer HSD test, P < 0.05).

**Supplemental Figure S7**. The effect of allantoin infiltration to the oldest leaves on its level in old, milled and young leaves of WT and *Atxdh1* (SALK_148366) mutant plants. The infiltration by injection of 5mM (A), 1mM (B) or 0 [control (H_2_O)] allantoin solutions into the oldest leaves (the first 4 leaves from the bottom) of 18 days old plants was performed by employing 1 ml needless syringe as previously described [see sulfite infiltration by injection (Brychkova *et al.*, 2012)]. Allantoin levels was determined 3 hour after the infiltration. The presented data is the mean obtained from 3 experiments. The values denoted with different letters are significantly different according to the Tukey-Kramer HSD test; p< 0.05. Different upper case letters indicate differences between mutants and WT plants.

**Supplemental Figure S8**. Immunoblot analysis of the large subunit (LSU) of Rubisco (A), D1, the component of the reaction center of PSII (B) and ATG8a, an autophagy protein (C). Proteins were extracted from old leaves (first 4 leaves counted from the bottom) of WT (Col) and *Atxdh1* mutant (SALK_148366) plants grown for 25 days on 1 or 5 mM nitrate as the only N source. 50 μg crude protein extracts was loaded into each lane.

**Supplemental Figure S9**. Leaf appearance (A), total chlorophyll content (B) and soluble protein content (C) of the old and young leaves of 18 days old WT (Col) and Atxdh1 (SALK_148366) mutant plants. Plants grew in nitrogen deficient soil supplemented with one-half strength Hoagland nutrient solutions containing 1 or 5mM NaNO_3_ as the only N source. The data represent the mean obtained from three independent experiments. Values denoted by different letters are significantly different (Tukey-Kramer HSD test, P < 0.05).

**Supplemental Tables**

**Supplemental Table S1**. Gene-specific primer sequences used for the expression analyses

## REFERENCES

Aerts R. 1990. Nutrient use efficiency in evergreen and deciduous species from heathlands. Oecologia 84, 391–397.

Agrimi G, Russo A, Pierri CL, Palmieri F. 2012. The peroxisomal NAD+ carrier of Arabidopsis thaliana transports coenzyme A and its derivatives. Journal of Bioenergetics and Biomembranes 44, 333–340.

Alamillo JM, Díaz-Leald J, Sánchez-Moran M, Pineda M. 2010. Molecular analysis of ureide accumulation under drought stress in Phaseolus vulgaris L. Plant, Cell and Environment 33, 1828–1837.

Amarante L do, Lima JD, Sodek L. 2006. Growth and stress conditions cause similar changes in xylem amino acids for different legume species. Environmental and Experimental Botany 58, 123–129.

Ashihara H, Sano H, Crozier A. 2008. Caffeine and related purine alkaloids: Biosynthesis, catabolism, function and genetic engineering. Phytochemistry 69, 841–856.

Atkins CA, Ritchie A, Rowe PB, McCairns E, Sauer D. 1982. De Novo Purine Synthesis in Nitrogen-Fixing Nodules of Cowpea (Vigna unguiculata [L.] Walp.) and Soybean (Glycine max [L.] Merr.). Plant Physiology 70, 55–60.

Bradford MM. 1976. A rapid and sensitive method for the quantitation of microgram quantities of protein using the principle of protein dye binding. Analytical Biochemistry 72, 248–254.

Brychkova G, Alikulov Z, Fluhr R, Sagi M. 2008. A critical role for ureides in dark and senescence-induced purine remobilization is unmasked in the Atxdh1 Arabidopsis mutant. Plant Journal 54, 496–509.

Brychkova G, Yarmolinsky D, Batushansky A, Grishkevich V, Khozin-Goldberg I, Fait A, Amir R, Fluhr R, Sagi M. 2015. Sulfite Oxidase Activity Is Essential for Normal Sulfur, Nitrogen and Carbon Metabolism in Tomato Leaves. Plants 4, 573–605.

Campbell WH. 1988. Nitrate reductase and its role in nitrate assimilation in plants. Physiologia Plantarum 74, 214–219.

Cataldo D a, Haroon M, Schrader LE, Youngs VL. 1975. Rapid colorimetric determination of nitrate in plant tissue by nitration of salicylic acid. Communications in Soil Science and Plant Analysis 6, 71–80.

Chalker-Scott L. 1999. Environmental significance of anthocyanins in plant stress responses. Photochemistry and Photobiology 70, 1–9.

Collier R, Tegeder M. 2012. Soybean ureide transporters play a critical role in nodule development, function and nitrogen export. Plant Journal 72, 355–367.

Coruzzi GM. 2003. Primary N-assimilation into Amino Acids in Arabidopsis. The Arabidopsis Book 2, 1–17.

Crafts Brandner SJ, Regina H, Urs F. 1998. Influence of nitrogen deficiency on senescence and the amounts of RNA and proteins in wheat leaves. Physiologia Plantarum 102, 192–200.

Crafts-Brandner SJ, Klein RR, Klein P, Hölzer R, Feller U. 1996. Coordination of protein and mRNA abundances of stromal enzymes and mRNA abundances of the Clp protease subunits during senescence of Phaseolus vulgaris (L.) leaves. Planta 200, 312–8.

Desimone M, Catoni E, Ludewig U, Hilpert M, Schneider A, Kunze R, Tegeder M, Frommer WB, Schumacher K. 2002. A novel superfamily of transporters for allantoin and other oxo derivatives of nitrogen heterocyclic compounds in Arabidopsis. The Plant cell 14, 847–856.

Diaz C, Purdy S, Christ A, Morot-Gaudry J-F, Wingler A. C. 2005. Characterization of markers to determine the extent and variability of leaf senescence in Arabidopsis. A metabolic profiling approach. Plant Physiology 138, 898–908.

Díaz-Leal J, Gálvez-Valdivieso G, Fernández J, Pineda M, Alamillo J. 2012. Developmental effects on ureide levels are mediated by tissue-specific regulation of allantoinase in Phaseolus vulgaris L. Journal of Experimental Botany 63, 695–709.

Eckhardt U, Grimm B, Hörtensteiner S. 2004. Recent advances in chlorophyll biosynthesis and breakdown in higher plants. Plant Molecular Biology 56, 1–14.

Fischer AM. 2007. Nutrient Remobilization During Leaf Senescence. Senescence Processes in Plants, 87–107.

Gepstein S, Sabehi G, Carp MJ, Hajouj T, Nesher MFO, Yariv I, Dor C, Bassani M. 2003. Large-scale identification of leaf senescence-associated genes. Plant Journal 36, 629–642.

Gould KS, Mckelvie J, Markham KR. 2002. Do anthocyanins function as antioxidants in leaves? Imaging of H2O2 in red and green leaves after mechanical injury. Journal of Experimental Botany 51, 123–129.

Hauck OK, Scharnberg J, Escobar NM, Wanner G, Giavalisco P, Witte C-P. 2014. Uric acid accumulation in an Arabidopsis urate oxidase mutant impairs seedling establishment by blocking peroxisome maintenance. The Plant cell 26, 3090–100.

Havé M, Marmagne A, Chardon F, Masclaux-Daubresse C. 2016. Nitrogen remobilisation during leaf senescence: lessons from Arabidopsis to crops. Journal of Experimental Botany 68, 2513–2529.

Hensel LL. 1993. Developmental and Age-Related Processes that influence the longevity and senescence of photosynthetic tissues in arabidopsis. The Plant cell 5, 553–564.

Hoagland DR, Arnon DI. 1950. The water-culture method for growing plants without soil. California Agricultural Experiment Station Circular 347, 1–32.

Hörtensteiner S, Feller U. 2002. Nitrogen metabolism and remobilization during senescence. Journal of experimental botany 53, 927–937.

Joy KW. 1988. Ammonia, glutamine, and asparagine: a carbon-nitrogen interface. Canadian Journal of Botany 66, 2103–2109.

Kaiser WM, Huber SC. 1994. Posttranslational Regulation of Nitrate Reductase in Higher Plants. Plant Physiology 106, 817–821.

Keren N, Berg A, Van Kan PJM, Levanon H, Ohad I. 1997. Mechanism of photosystem II photoinactivation and D1 protein degradation at low light: The role of back electron flow. Plant Biology 94, 1579–1584.

Kojima S, Bohner A, Gassert B, Yuan L, Wirén N Von. 2007. AtDUR3 represents the major transporter for high-affinity urea transport across the plasma membrane of nitrogen-deficient Arabidopsis roots. Plant Journal 52, 30–40.

Koyama Y, Tomoda Y, Kato M, Ashihara H. 2003. Metabolism of purine bases, nucleosides and alkaloids in theobromine-forming Theobroma cacao leaves. Plant Physiology and Biochemistry 41, 977–984.

Krapp A, Berthome R, Orsel M, Mercey-Boutet S, Yu A, Castaings L, Elftieh S, Major H, Renou JP, Daniel-Vedele F. 2011. Arabidopsis Roots and Shoots Show Distinct Temporal Adaptation Patterns toward Nitrogen Starvation. Plant Physiology 157, 1255–1282.

Laby RJ, Kincaid MS, Kim D, Gibson SI. 2000. The Arabidopsis sugar-insensitive mutants sis4 and sis5 are defective in abscisic acid synthesis and response. Plant Journal 23, 587–596.

Lange PR, Geserick C, Tischendorf G, Zrenner R. 2007. Functions of Chloroplastic Adenylate Kinases in Arabidopsis. Plant Physiology 146, 492–504.

Lescano CI, Martini C, González CA, Desimone M. 2016. Allantoin accumulation mediated by allantoinase downregulation and transport by Ureide Permease 5 confers salt stress tolerance to Arabidopsis plants. Plant Molecular Biology 91, 581–595.

Lim PO, Kim HJ, Nam HG. 2007. Leaf senescence. Annual Review of Plant Biology 58, 115–136.

Lim PO, Woo HR, Nam HG. 2003. Molecular genetics of leaf senescence in Arabidopsis. Trends in Plant Science 8, 272–278.

Liu J, Wu YH, Yang JJ, Liu YD, Shen FF. 2008. Protein degradation and nitrogen remobilization during leaf senescence. Journal of Plant Biology 51, 11–19.

Ma X, Wang W-M, Bittner F, et al. 2016. Dual and Opposing Roles of Xanthine Dehydrogenase in Defense-Associated Reactive Oxygen Species Metabolism in Arabidopsis. The Plant Cell 28, 1108–1126.

Masclaux C, Valadier MH, Brugière N, Morot-Gaudry JF, Hirel B. 2000. Characterization of the sink/source transition in tobacco ( Nicotiana tabacum L.) shoots in relation to nitrogen management and leaf senescence. Planta 211, 510–8.

Masclaux-Daubresse C, Daniel-Vedele F, Dechorgnat J, Chardon F, Gaufichon L, Suzuki A. 2010. Nitrogen uptake, assimilation and remobilization in plants: Challenges for sustainable and productive agriculture. Annals of Botany 105, 1141–1157.

Masclaux-Daubresse C, Reisdorf-Cren M, Orsel M. 2008. Leaf nitrogen remobilisation for plant development and grain filling. Plant Biology 10, 23–36.

Matile P, Hortensteiner S, Thomas H, Krautler B. 1996. Chlorophyll Breakdown in Senescent Leaves. Plant Physiology 112, 1403–1409.

Meyer R, Wagner KG. 1986. Nucleotide pools in leaf and root-tissue of tobacco plants: Influence of leaf senescence. Physiologia Plantarum 67, 666–672.

Miller JD, Arteca RN, Pell EJ. 1999. Senescence-associated gene expression during ozone-induced leaf senescence in Arabidopsis. Plant Physiology 120, 1015–24.

Molins-Legua C, Meseguer-Lloret S, Moliner-Martinez Y, Campíns-Falcó P. 2006. A guide for selecting the most appropriate method for ammonium determination in water analysis. TrAC - Trends in Analytical Chemistry 25, 282–290.

Munné-Bosch S, Alegre L. 2004. Die and let live: Leaf senescence contributes to plant survival under drought stress. Functional Plant Biology 31, 203–216.

Murashige T, Skoog F. 1962. A revised medium for rapid growth and bioassays with tobacco tissue cultures. Physiologia plantarum 15, 473–497.

Murray JD, Liu C-W, Chen Y, Miller AJ. 2016. Nitrogen sensing in legumes. Journal of Experimental Botany 68, 1919–1926.

Nakagawa A, Sakamoto S, Takahashi M, Morikawa H, Sakamoto A. 2007. The RNAi-mediated silencing of xanthine dehydrogenase impairs growth and fertility and accelerates leaf senescence in transgenic Arabidopsis plants. Plant and Cell Physiology 48, 1484–95.

Pageau K, Reisdorf-Cren M, Morot-Gaudry JF, Masclaux-Daubresse C. 2006. The two senescence-related markers, GS1 (cytosolic glutamine synthetase) and GDH (glutamate dehydrogenase), involved in nitrogen mobilization, are differentially regulated during pathogen attack and by stress hormones and reactive oxygen species in Nicoti. Journal of Experimental Botany 57, 547–557.

Park S-Y, Yu J-W, Park J-S, et al. 2007. The senescence-induced staygreen protein regulates chlorophyll degradation. The Plant cell 19, 1649–64.

Pate JS. 1980. Transport and Partitioning of Nitrogenous Solutes. Annual Review of Plant Physiology 31, 313–340.

Pruzinská, A, Tanner G, Aubry S, et al. 2005. Chlorophyll Breakdown in Senescent Arabidopsis Leaves. Characterization of Chlorophyll Catabolites and of Chlorophyll Catabolic Enzymes Involved in the Degreening Reaction 1. Plant Physiology 139, 52–63.

Reumann S, Babujee L, Ma C, Wienkoop S, Siemsen T, Antonicelli GE, Rasche N, Lüder F, Weckwerth W, Jahn O. 2007. Proteome analysis of Arabidopsis leaf peroxisomes reveals novel targeting peptides, metabolic pathways, and defense mechanisms. The Plant cell 19, 3170–3193.

Ritchie RJ. 2006. Consistent sets of spectrophotometric chlorophyll equations for acetone, methanol and ethanol solvents. Photosynthesis Research 89, 27–41.

Rothe GM. 1974. Aldehyde oxidase isoenzymes (E.C. 1. 2. 3.1) in potato tubers (Solanum tuberosum). Plant and Cell Physiology 499, 493–499.

Sabina RL, Paul A-L, Ferl RJ, Laber B, Lindell SD. 2007. Adenine nucleotide pool perturbation is a metabolic trigger for AMP deaminase inhibitor-based herbicide toxicity. Plant Physiology 143, 1752–1760.

Sagi M, Fluhr R. 2001. Superoxide production by plant homologues of the gp91(phox) NADPH oxidase. Modulation of activity by calcium and by tobacco mosaic virus infection. Plant Physiol 126, 1281–1290.

Sagi M, Lips HS. 1998. The levels of nitrate reductase and MoCo in annual ryegrass as affected by nitrate and ammonium nutrition. Plant Science 135, 17–24.

Sagi M, Omarov RT, Lips SH. 1998. The Mo-hydroxylases xanthine dehydrogenase and aldehyde oxidase in ryegrass as affected by nitrogen and salinity. Plant Science 135, 125–135.

Sagi M, Savidov NA, Lvov NP, Lips SH. 1997. Nitrate reductase and molybdenum cofactor in annual ryegrass as affected by salinity and nitrogen source. Physiologia Plantarum 99, 546–553.

Schmidt A, Baumann N, Schwarzkopf A, Frommer WB, Desimone M. 2006. Comparative studies on Ureide Permeases in Arabidopsis thaliana and analysis of two alternative splice variants of AtUPS5. Planta 224, 1329–1340.

Schmidt S, Stewart GR. 1998. Transport, storage and mobilization of nitrogen by trees and shrubs in the wet/dry tropics of northern Australia. Tree Physiology 18, 403–410.

Schmidt A, Su YH, Kunze R, Warner S, Hewitt M, Slocum RD, Ludewig U, Frommer WB, Desimone M. 2004. UPS1 and UPS2 from Arabidopsis mediate high affinity transport of uracil and 5-fluorouracil. Journal of Biological Chemistry 279, 44817–44824.

Schroeder RY, Zhu A, Eubel H, Dahncke K, Witte C-P. 2017. The ribokinases of Arabidopsis thaliana and Saccharomyces cerevisiae are required for ribose recycling from nucleotide catabolism, which in plants is not essential to survive prolonged dark stress. New Phytologist, nph.14782.

Schubert KR. 1981. Enzymes of Purine Biosynthesis and Catabolism in Glycine max: I. comparison of activities with N(2) fixation and composition of xylem exudate during nodule development. Plant Physiology 68, 1115–1122.

Schubert KR. 1986. Products of biological nitrogen fixation in higher plants: synthesis, transport, and metabolism. Annual Review of Plant Physiology 37, 539–574.

Schussler MD, Alexandersson E, Bienert GP, Kichey T, Laursen KH, Johanson U, Kjellbom P, Schjoerring JK, Jahn TP. 2008. The effects of the loss of TIP1;1 and TIP1;2 aquaporins in Arabidopsis thaliana. Plant Journal 56, 756–767.

Simpson RJ, Lambers H, Dalling MJ. 1983. Nitrogen Redistribution during Grain Growth in Wheat (Triticum aestivum L.): IV. Development of a Quantitative Model of the Translocation of Nitrogen to the Grain. Plant Physiology 71, 7–14.

Sivasankar S, Oaks A. 1996. Nitrate assimilation in higher plants: The effect of metabolites and light. Plant Physiology and Biochemistry 34, 609–620.

Smith PMC, Atkins CA. 2002. Purine Biosynthesis. Big in Cell Division, Even Bigger in Nitrogen Assimilation. Plant Physiology 128, 793–802.

Somerville CR, Ogren WL. 1980. Inhibition of photosynthesis in Arabidopsis mutants lacking leaf glutamate synthase activity. Nature 286, 257–259.

Srivastava S, Brychkova G, Yarmolinsky D, Soltabayeva A, Samani T, Sagi M. 2017. Aldehyde Oxidase 4 plays a critical role in delaying silique senescence by catalyzing aldehyde detoxification. Plant Physiology, pp.01939.2016.

Staswick P. 1997. The occurrence and gene expression of vegetative storage proteins and a Rubisco Complex Protein in several perennial soybean species. Journal of Experimental Botany 48, 2031–2036.

Stebbins NE, Polacco JC. 1995. Urease is not essential for ureide degradation in soybean. Plant Physiology 109, 169–175.

Suzuki Y, Shioi Y. 1999. Detection of Chlorophyll Breakdown Products in the Senescent Leaves of Higher Plants. Plant and Cell Physiology 40, 909–915.

Ta CT, Weiland RT. 1992. Nitrogen Partitioning in Maize during Ear Development. Crop Science 32, 443.

Takagi H, Ishiga Y, Watanabe S, et al. 2016. Allantoin, a stress-related purine metabolite, can activate jasmonate signaling in a MYC2-regulated and abscisic acid-dependent manner. Journal of Experimental Botany 67, 2519–2532.

Tanaka R, Hirashima M, Satoh S, Tanaka A. 2003. The Arabidopsis-accelerated cell death Gene ACD1 is Involved in Oxygenation of Pheophorbide a: Inhibition of the Pheophorbide a Oxygenase Activity does not Lead to the “Stay-Green” Phenotype in Arabidopsis. Plant and Cell Physiology 44, 1266–1274.

Thomas RJ, Feller U, Erismann KH. 1980. Ureide metabolism in non-nodulated Phaseolus vulgaris L. Journal of Experimental Botany 31, 409–417.

Todd CD, Polacco JC. 2004. Soybean cultivars “Williams 82” and “Maple Arrow” produce both urea and ammonia during ureide degradation. Journal of Experimental Botany 55, 867–877.

Todd CD, Polacco JC. 2006. AtAAH encodes a protein with allantoate amidohydrolase activity from Arabidopsis thaliana. Planta 223, 1108–1113.

Todd CD, Tipton PA, Blevins DG, Piedras P, Pineda M, Polacco JC. 2006. Update on ureide degradation in legumes. Journal of Experimental Botany 57, 5–12.

Uhart SA, Andrade FH. 1995. Nitrogen and carbon accumulation and remobilization during grain filling in maize under different source/sink ratios. Crop Science 35, 183–190.

Vogels GD, Van Der Drift C. 1970. Differential analyses of glyoxylate derivatives. Analytical Biochemistry 33, 143–157.

Watanabe S, Matsumoto M, Hakomori Y, Takagi H, Shimada H, Sakamoto A. 2014. The purine metabolite allantoin enhances abiotic stress tolerance through synergistic activation of abscisic acid metabolism. Plant, Cell and Environment 37, 1022–1036.

Wendler R, Carvalho PO, Pereira JS, Millard P. 1995. Role of nitrogen remobilization from old leaves for new leaf growth of Eucalyptus globulus seedlings. Tree Physiology 15, 679–83.

Werner, AK, Medina-Escobar N, Zulawski M, Sparkes, I a., Cao F-Q, Witte, C-P. 2013. The Ureide-Degrading Reactions of Purine Ring Catabolism Employ Three Amidohydrolases and One Aminohydrolase in Arabidopsis, Soybean, and Rice. Plant Physiology 163, 672–681.

Werner a. K, Romeis T, Witte C-P. 2010. Ureide catabolism in Arabidopsis thaliana and Escherichia coli. Nature Chemical Biology 6, 19–21.

Werner, a. K, Sparkes, I a., Romeis T, Witte C-P. 2008. Identification, Biochemical Characterization, and Subcellular Localization of Allantoate Amidohydrolases from Arabidopsis and Soybean. Plant Physiology 146, 418–430.

Werner AK, Witte CP. 2011. The biochemistry of nitrogen mobilization: Purine ring catabolism. Trends in Plant Science 16, 381–387.

Xu J, Zhang HY, Xie CH, Xue HW, Dijkhuis P, Liu CM. 2005. EMBRYONIC FACTOR 1 encodes an AMP deaminase and is essential for the zygote to embryo transition in Arabidopsis. Plant Journal 42, 743–756.

Yeoh HH, Wee YC. 1994. Leaf protein contents and nitrogen-to-protein conversion factors for 90 plant species. Food Chemistry 49, 245–250.

Yesbergenova Z, Yang G, Oron E, Soffer D, Fluhr R, Sagi M. 2005. The plant Mo-hydroxylases aldehyde oxidase and xanthine dehydrogenase have distinct reactive oxygen species signatures and are induced by drought and abscisic acid. Plant Journal 42, 862–876.

Yoshino M, Murakami K, Tsushima K. 1979. AMP deaminase from baker’s yeast. Purification and some regulatory properties. BBA - Enzymology 570, 157–166.

Zrenner R, Stitt M, Sonnewald U, Boldt R. 2006. Pyrimidine and purine biosynthesis and degradation in plants. Annual review of plant biology 57, 805–836.

